# Systematic evaluation of robustness of deconvolution methods for spatial transcriptomics data in case of cell type mismatch

**DOI:** 10.1101/2025.08.12.669903

**Authors:** Utkarsh M. Mahamune, Aldo Jongejan, Antoine H. C. van Kampen, Lisa G. M. van Baarsen, Perry D. Moerland

## Abstract

Sequencing-based spatial transcriptomics (ST) approaches preserve spatial information but with limited cellular resolution, whereas single-cell RNA-sequencing (scRNA-seq) techniques provide single-cell resolution but lose spatial context during tissue dissociation. Given these complementary strengths, computational tools have been developed to combine scRNA-seq and ST data. These methods use deconvolution techniques to identify cell types and estimate their proportions at each spatial location in ST data, using scRNA-seq reference data. However, these methods are sensitive to missing cell types in the scRNA-seq reference, a problem known as cell type mismatch. Using two reference datasets, we performed extensive simulations to systematically evaluate the robustness to cell type mismatch of six deconvolution methods (CARD, cell2location, RCTD, Seurat, SPOTlight, Stereoscope) tailored for ST data, and two designed for bulk RNA-seq data (MuSiC, SCDC). At baseline, that is, with no cell types missing from the reference datasets, cell2location showed the strongest performance, while Seurat performed the worst. By simulating different cell type mismatch scenarios, we found that the performance of deconvolution methods decreases proportionally to the number of cell types missing from the reference. Moreover, compared to baseline, for most methods the relative decrease in performance is similar. Additionally, methods that perform well at baseline tend to assign the proportions of a missing cell type to the transcriptionally most similar cell types present in the reference data.

Our results highlight the adverse effects of cell type mismatch on the performance of deconvolution methods for ST data and stress the need for more robust approaches to this issue.

## 1. Introduction

In the past decade, single-cell RNA sequencing (scRNA-seq) methods have developed immensely. Current scRNA-seq techniques can capture and characterize the transcriptional profiles of thousands of cells at single-cell resolution [1]. This development has paved the way for exciting opportunities to study cellular heterogeneity. However, for tissue-based analyses, the spatial context of these cells is lost during the isolation of single cells before sequencing. Recently, spatially resolved transcriptomics techniques have been developed to study intact tissue, while preserving spatial information. These spatial transcriptomics (ST) approaches can be divided into two main categories. The first category comprises imaging-based spatial transcriptomics [2, 3], which achieves single-cell resolution but for a limited set of transcripts only. The second category consists of sequencing-based techniques [4-7] that generate transcriptome-wide gene expression profiles but are unable to achieve single-cell resolution. Consequently, these techniques provide cumulative gene expression levels of a few to dozens of cells present at a specific spatial location (hereinafter referred to as a ‘spot’). Newer sequencing-based ST techniques increase the resolution to near single-cell level [6, 7] or even lower [5, 8], but these are still limited by the number of genes that can be detected. Here, we focus on sequencing-based ST techniques, as these are most commonly used in biomedical life science research.

Due to their limited resolution, the spot-level measurements of sequencing-based techniques may represent the expression of multiple cells of different cell types. A fundamental challenge is therefore to determine the proportions of the different cell types present in such a spot. A large number of deconvolution methods have been developed for this purpose. One class of approaches consists of reference-free deconvolution methods, which use only the ST data [9-13]. The other class of approaches are reference-based deconvolution methods, which use scRNA-seq reference data along with ST data to estimate cell-type proportions within each spot in ST data. In this context, the reference data consist of high-resolution single-cell gene expression profiles that provide representative expression signatures for each cell type. Reference-based deconvolution methods have already been used for many years on bulk RNA-seq data [14-18]. Methods developed for bulk RNA-seq data can, in principle, be applied to ST data too. However, dedicated methods that take into account specific characteristics of ST data [10, 19-25], such as a limited number of cells and cell types in a spot, have been shown to often outperform methods developed for bulk RNA-seq data [19, 21, 25]. Comprehensive benchmarking studies have also shown that reference-based methods in general outperform reference-free methods [26-28]. Moreover, reference-free methods still rely on external data to annotate identified topics [11] or metagenes [12] using known cell type markers. Here, we focus exclusively on reference-based deconvolution methods.

Given the dependence of reference-based methods on matching scRNA-seq data, it has been suggested that these methods are sensitive to the absence of cell types in either the reference or ST data [29], a problem referred to as cell type mismatch. Ideally, both datasets include the same cell types for optimal deconvolution results. However, in practice this is often not the case, because of differences in the experimental protocols and measurement characteristics [29]. For example, certain cell types may be depleted or missing in scRNA-seq data as a result of the tissue dissociation procedure. Cell type mismatch can also occur if atlas-based reference data is used in which a cell type of interest is missing or because of the sorting strategy applied for the creation of the scRNA-seq reference data [30].

In several recent studies [26-28, 31], the performance of deconvolution approaches for ST data has been comprehensively and independently benchmarked to investigate the robustness of deconvolution methods to sequencing depth and spot size, as well as to evaluate the accuracy and usability of the method. However, these studies did not systematically assess the robustness of deconvolution methods to cell type mismatch. Robustness to cell type mismatch has only been assessed for a specific example with one or two cell types missing from the reference data for the RCTD deconvolution method [20]. Not surprisingly, these results indicated reduced performance of RCTD in case of cell type mismatch, especially when no transcriptionally similar cell type is present in the reference data. Robustness of CARD and several other deconvolution methods has also been evaluated, but only for scenarios with one cell type missing from the reference [10]. In this case too, cell type mismatch led to reduced performance, with CARD being more robust than the other methods. To our knowledge, no systematic evaluation has yet assessed the robustness of deconvolution methods to cell type mismatch. This is crucial because, in practice, some cell types are frequently underrepresented or lost during tissue dissociation, resulting in incomplete reference data.

Here, we go beyond previous work, which addressed cell type mismatch only in isolated cases or for individual methods, by systematically quantifying robustness trends as a function of mismatch severity across multiple ST deconvolution methods, reference datasets, and simulated ST datasets. For this purpose, we use extensive simulations to systematically evaluate the robustness to cell type mismatch of eight deconvolution methods. We present an algorithm for simulating ST data, enabling us to vary essential spatial data characteristics. The methods evaluated consist of six widely used state-of-the-art reference-based deconvolution methods designed for ST data (CARD [10], cell2location [24], RCTD [20], Seurat [22], SPOTlight [21], and Stereoscope [19]) and two bulk RNA-seq-specific methods (MuSiC [17], SCDC [18]). We also examine how each deconvolution method redistributes the proportions of a missing cell type among the remaining cell types in the reference datasets. We show that deconvolution performance decreases proportionally to the number of cell types missing from the reference data and that the relative decrease in performance is similar across most deconvolution methods. This highlights the need for deconvolution methods for ST data that are more robust to cell type mismatch.

## 2. Materials and methods

Detailed information can be found in the Supplementary Information.

### 2.1 Single-cell reference data

Lymph node single-cell reference. We used two human lymph node scRNA-seq datasets to construct an integrated reference dataset. The Tabula Sapiens Consortium lymph node data (version 3) [32] comprises 53,275 cells, most of which are B or T cells. Since this dataset contains only few fibroblasts, we also included a lymph node stromal cell (LNSC) dataset consisting of 13,850 cells [33]. The datasets were further processed and integrated using Seurat (v4.1.0) [22]. Preprocessing steps, including quality control thresholds, SingleR [34] reannotation, integration, gene selection, and visualization, are described in Supplementary Information S1.1-S1.3.

After integration using Seurat’s anchor-based workflow, we retained 13 cell types with sufficient representation and downsampled overrepresented cell types to 300 cells each to create a balanced reference. This resulted in 3,519 cells across 13 annotated cell types. Gene filtering following the strategy described by Kleshchevnikov et al. [24], yielded 11,474 informative genes. Filtering criteria and gene selection rules are described in Supplementary Information S1.4-S1.5.

This integrated dataset served both as the single-cell reference for all cell type deconvolution methods and as the basis for generating simulated ST data.

Hypothalamus single-nucleus reference. In addition, we used a hypothalamus single-nucleus dataset from the Human Brain Cell Atlas as an independent validation set. The characteristics of this dataset and processing steps are described in detail in Supplementary Information S5.

### 2.2 Cell type deconvolution methods

In our analysis, we benchmarked eight reference-based deconvolution methods: six designed specifically for spatial transcriptomics (CARD [10], cell2location [24], RCTD [20], Seurat [22], SPOTlight [21], Stereoscope [19]) and two originally developed for bulk RNA-seq data (MuSiC [17], SCDC [18]).

These methods represent a range of modelling strategies: non-negative matrix factorization (NMF; CARD), hierarchical Bayesian modelling (cell2location), Poisson-based likelihood estimation (RCTD), label transfer through canonical correlation analysis (Seurat), NMF combined with non-negative least squares (SPOTlight), and probabilistic modelling with negative binomial distributions (Stereoscope). MuSiC and SCDC both use weighted non-negative least squares frameworks, treating each spatial spot as an independent bulk RNA-seq sample. All methods were run according to their recommended workflows, with only minor adjustments to accommodate the structure of the simulated datasets. Full details on parameter settings, software versions, and preprocessing steps are provided in Supplementary Information S2.

### 2.3 Generation of simulated spatial transcriptomics data

To evaluate deconvolution accuracy under controlled conditions, we developed a simulation procedure that generates ST datasets with known ground-truth cell type proportions (Figure S2). The simulations draw directly from the reference datasets described above.

Each simulated dataset consisted of 1,600 spatial spots arranged on a 40 × 40 grid. For each spot, we sampled:

1. Number of cell types present,
2. Which cell types appear,
3. Proportions of each cell type,
4. Total number of cells, and
5. Cell-type-specific mRNA profiles based on averages of five randomly selected reference cells.

Counts for each spot were generated by summing contributions from all underlying cell types and downsampled to approximate 10x Visium sequencing depth. Note that since the selected deconvolution methods, with the exception of CARD, do not take spatial correlation into account, spots are treated as independent in our simulation framework. Exact mathematical definitions, sampling distributions, the multinomial allocation, and per-gene count calculations are documented in Supplementary Information S3.

Using the lymph node reference data, we generated three simulated datasets with varying levels of cellular complexity, which were used for downstream evaluations:

– ST1: 4–8 cell types, 10–15 cells per spot
– ST2: 1–5 cell types, 10–15 cells per spot
– ST3: 1–5 cell types, 3–7 cells per spot (number of cells ≥ number of cell types)

For the hypothalamus reference data, we only generated ST1.

### 2.4 Benchmark framework and performance evaluation

Figure 1 summarizes our benchmarking approach. The framework consisted of two major components: (1) evaluating baseline performance when all cell types are present in the reference data, and (2) assessing robustness when reference cell types are systematically removed.

**Figure 1.**
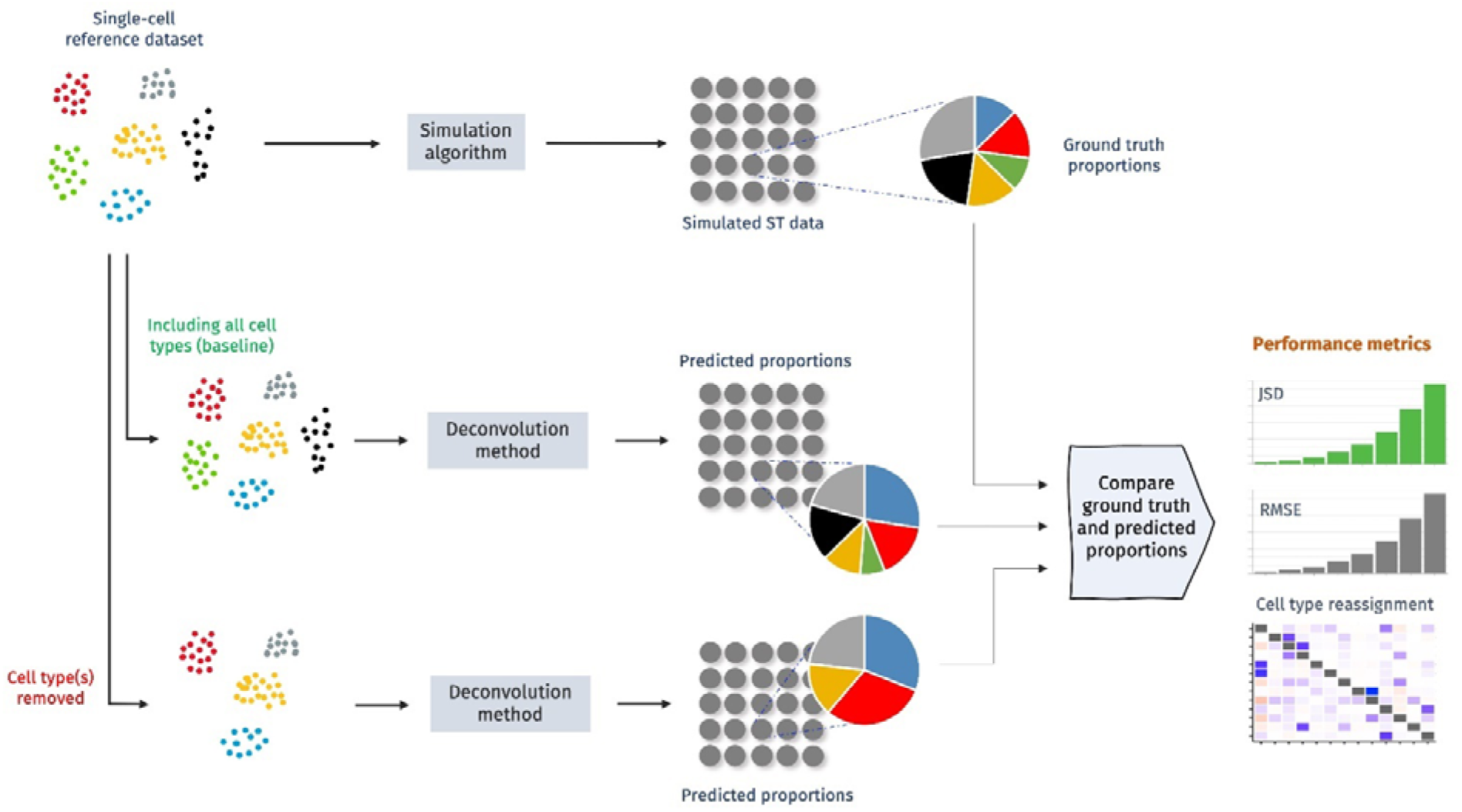
Benchmark framework. A scRNA-seq dataset is used both as a reference dataset and as input for simulating spatial transcriptomics (ST) data with known per spot cell type proportions (ground truth). Selected reference-based deconvolution methods are then applied in the baseline scenario, where both the single-cell reference and the simulated ST data contain the same cell types. Robustness to cell type mismatch is assessed by systematically removing one or more cell types from the reference data, but not from the ST data, and applying the same deconvolution methods. For both scenarios, we compare the ground truth and predicted proportions using the Jensen-Shannon divergence (JSD) and the root mean square error (RMSE) as performance metrics. In the cell type mismatch scenario, we also determine the cell type reassignment values to assess to which cell type(s) the proportions of the removed cell type(s) are reassigned. The benchmark framework isolates the effect of cell type mismatch by comparing baseline and mismatch scenarios against known ground-truth cell type proportions.

#### Cell type mismatch scenarios

We simulated mismatch situations by removing reference cell types while keeping ST data unchanged. Because scRNA-seq reference datasets often have missing or depleted populations due to dissociation inefficiencies or experimental bias, this design reflects real-world conditions. We examined:

– Single cell type removal: each of the 13 (lymph node) or 14 (hypothalamus) cell types was removed individually,
– Pair and triplet removal: sequential removal of pairs or triplets of transcriptionally similar cell types,
– Quintuplet removal (lymph node only): removal of one predefined set consisting of five cell types,
– Extreme mismatch: sequential removal of 10 or 11 (lymph node) or 11 or 12 (hypothalamus) cell types, leaving only three or two cell types in the reference, respectively.

Exact definitions of pairs, triplets, and quintuplets, and the rationale based on expression similarity are given in Supplementary Information S4 and Table S2 for the lymph node reference data and in Supplementary Information S5 and Table S4 for the hypothalamus reference data.

#### Performance metrics

To compare predicted and ground-truth proportions, we used two widely applied measures:

– Jensen–Shannon divergence (JSD): a measure of similarity between predicted and true probability distributions (range 0–1; lower is better).
– Root mean square error (RMSE): quantifies mean prediction deviation across all cell types per spot (lower is better).

Equations of both metrics are provided in Supplementary Information S4.

#### Cell type reassignment analysis

To understand how individual methods redistribute the proportions of missing cell types relative to the baseline scenario, we defined the *reassignment value*. This value quantifies how much of the removed cell types’ proportions is shifted to each remaining cell type. A value close to 1 for a remaining cell type indicates that the removed cell type’s proportions are reassigned primarily to that cell type; values below 0 or above 1 reflect broader redistribution effects. The exact definition, together with pseudocode and numerical examples, are documented in Supplementary Information S4.

## 3. Results

We used lymph node single-cell data and hypothalamus single-nucleus data as reference datasets. The lymph node reference data is used to illustrate the main results, whereas the hypothalamus reference data is used as an independent validation dataset.

### 3.1 Cell type deconvolution methods show large differences in baseline performance

Using the integrated human lymph node dataset as the single-cell reference and as input for simulating ST data, we evaluated the performance of eight different cell type deconvolution methods. We simulated three ST datasets from the lymph node reference data varying both the number of cells and the number of cell types present per spot. Figure 2 shows the deconvolution method performance using the first simulated dataset (ST1), which on average contains more cells and/or cell types per spot than the other two datasets (ST2, ST3). Results for the latter datasets are shown in the Supplementary Information.

**Figure 2.**
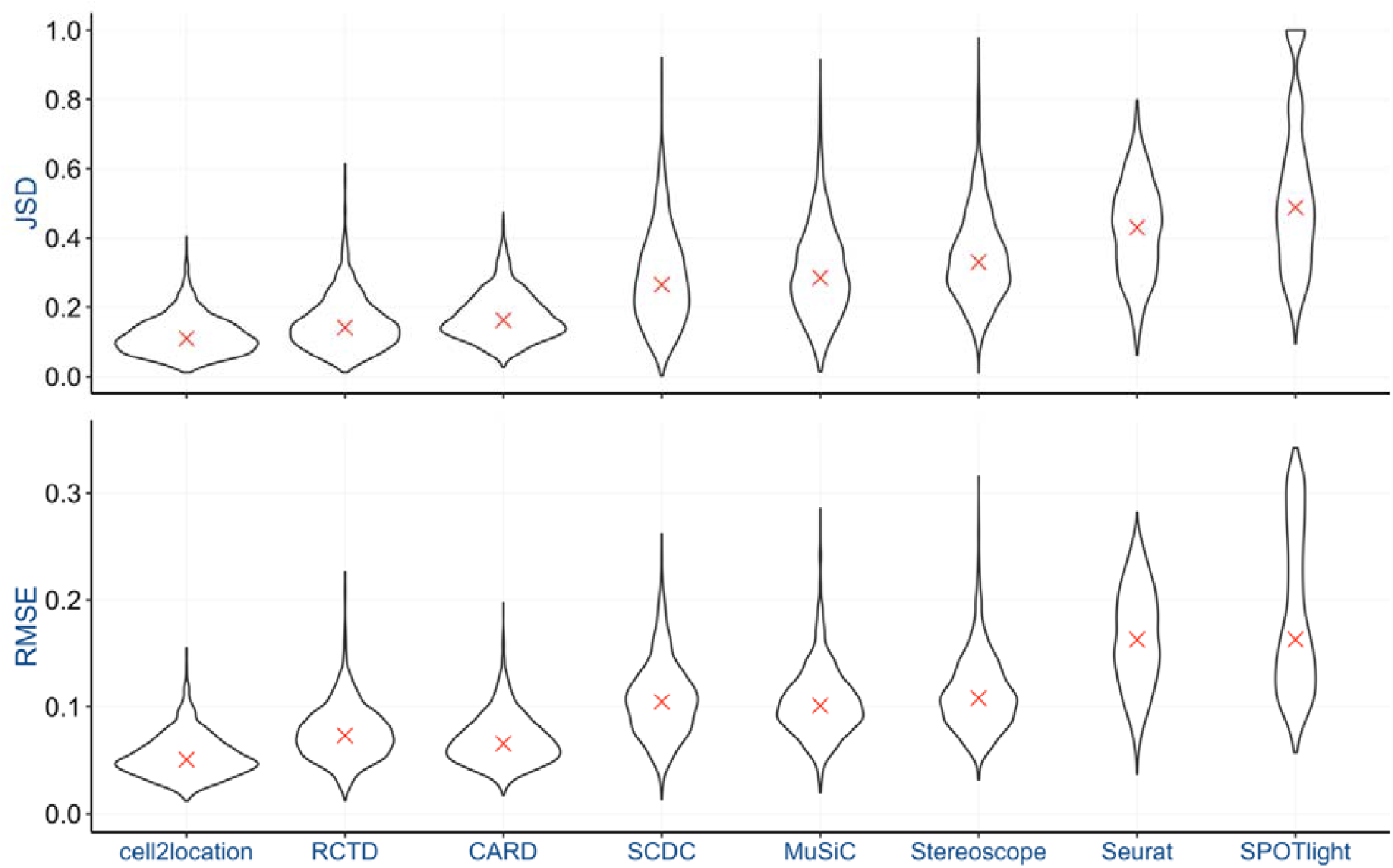
Baseline scenario performance of cell type deconvolution methods for the lymph node data. Comparison of the ground truth and predicted proportions using the Jensen-Shannon divergence (JSD) and the root mean square error (RMSE) as performance metrics for simulated dataset ST1 for eight cell type deconvolution methods. Violin plots correspond to the distribution of the values of the indicated performance metric across 1,600 spots. Median values are indicated with a red cross. Deconvolution methods in both panels are ordered by increasing median JSD value, with higher values indicating worse performance. Cell2location, RCTD, and CARD are most accurate at baseline, whereas Seurat, Stereoscope, and SPOTlight perform substantially worse.

First, we assessed the performance of these methods for the baseline scenario, when both the single-cell reference and the simulated ST data contain the same 13 cell types (Figure 1). We observed substantial performance differences between the deconvolution methods across both JSD and RMSE metrics (Figure 2). The top-performing methods are three spatial transcriptomics-specific cell type deconvolution methods (cell2location, RCTD, and CARD) with median JSD values of 0.11, 0.14 and 0.16, respectively, and median RMSE values of 0.05, 0.07 and 0.07, respectively. The two bulk RNA-seq-specific methods (SCDC, MuSiC) perform worse with median JSD values of 0.27 and 0.29, respectively, and median RMSE values of 0.10. The worst performing methods are Stereoscope, Seurat, and SPOTlight with median JSD values of 0.33, 0.43 and 0.49, respectively, and median RMSE values of 0.11, 0.16 and 0.16, respectively. Results for the other two simulated ST datasets were very similar in terms of the ranking on median JSD and RMSE values (Figure S3a, S3c).

To further characterize baseline performance, we also evaluated per-cell-type RMSE across the 13 lymph node cell types (Figure S4). The per-cell-type results are consistent with the method-level trends, with cell2location, RCTD, and CARD showing the lowest error across most cell types. Some cell types, such as myocytes and skeletal muscle cells, appeared more challenging across methods than others. Interestingly, the low accuracy of SPOTlight appears to be mainly driven by incorrect estimation of the fibroblast proportions. These results indicate that deconvolution accuracy reflects both overall method performance and cell-type-specific difficulty.

### 3.2 Performance decreases proportionally to the number of cell types missing from the reference data

At baseline no cell types are missing from the reference data. In order to evaluate the robustness to cell type mismatch, we systematically removed one or more cell types from the reference data, but not from the ST data, and then applied the selected deconvolution methods (Figure 1). We investigated cell type mismatch scenarios with 1, 2, 3, 5, 10 or 11 missing cell types from the 13 cell types in total. In case of one missing cell type, all 13 possibilities of removing a single cell type were evaluated. In case of 2, 3 or 5 missing cell types, we evaluated a limited number of combinations by systematically defining sets of transcriptionally similar cell types to be removed from the reference data (see Supplementary Information S4). We also evaluated two extreme scenarios with 10 or 11 missing cell types, where only 3 or 2 cell types remain, respectively. In this case, we only included the triplets or pairs of transcriptionally similar cell types defined above in the reference data. For all these scenarios we determined the spot-wise JSD and RMSE and used these to calculate the spot-wise difference in performance (ΔJSD, ΔRMSE) between a scenario with cell types missing from the reference data and the baseline scenario (Figure 2).

Both in terms of ΔJSD and ΔRMSE, the cell type mismatch results show a clear trend. Performance decreases proportionally to the number of cell types missing compared to the baseline scenario (Figure 3, Figure S5a). This trend is highly consistent for the methods that performed best at baseline (cell2location, RCTD, CARD). Specifically, this means that the relative ranking of the cell type deconvolution methods in the case of cell type mismatch is largely determined by their performance at baseline. The most straightforward explanation is that, as more cell types are removed from the reference data, a larger fraction of the signal must be redistributed across remaining cell types, causing performance to decrease proportionally to the number of missing cell types. For the worst performing methods at baseline (Stereoscope, Seurat, SPOTlight), the decrease in performance is less pronounced. This can be explained by the fact that performance at baseline is already low and therefore cannot decrease much more. Note that CARD failed to execute with only two cell types present in the reference data. Furthermore, Seurat was unable to find anchors between the single-cell reference and ST data with either two or three cell types across certain pairs and triplets. As a result, it failed to perform in those cases, so we indicated the results as missing for that scenario. Results for the other two simulated ST datasets showed similar trends in terms of the ΔJSD and ΔRMSE (Figure S3b, S3d, S5b, S5c).

**Figure 3.**
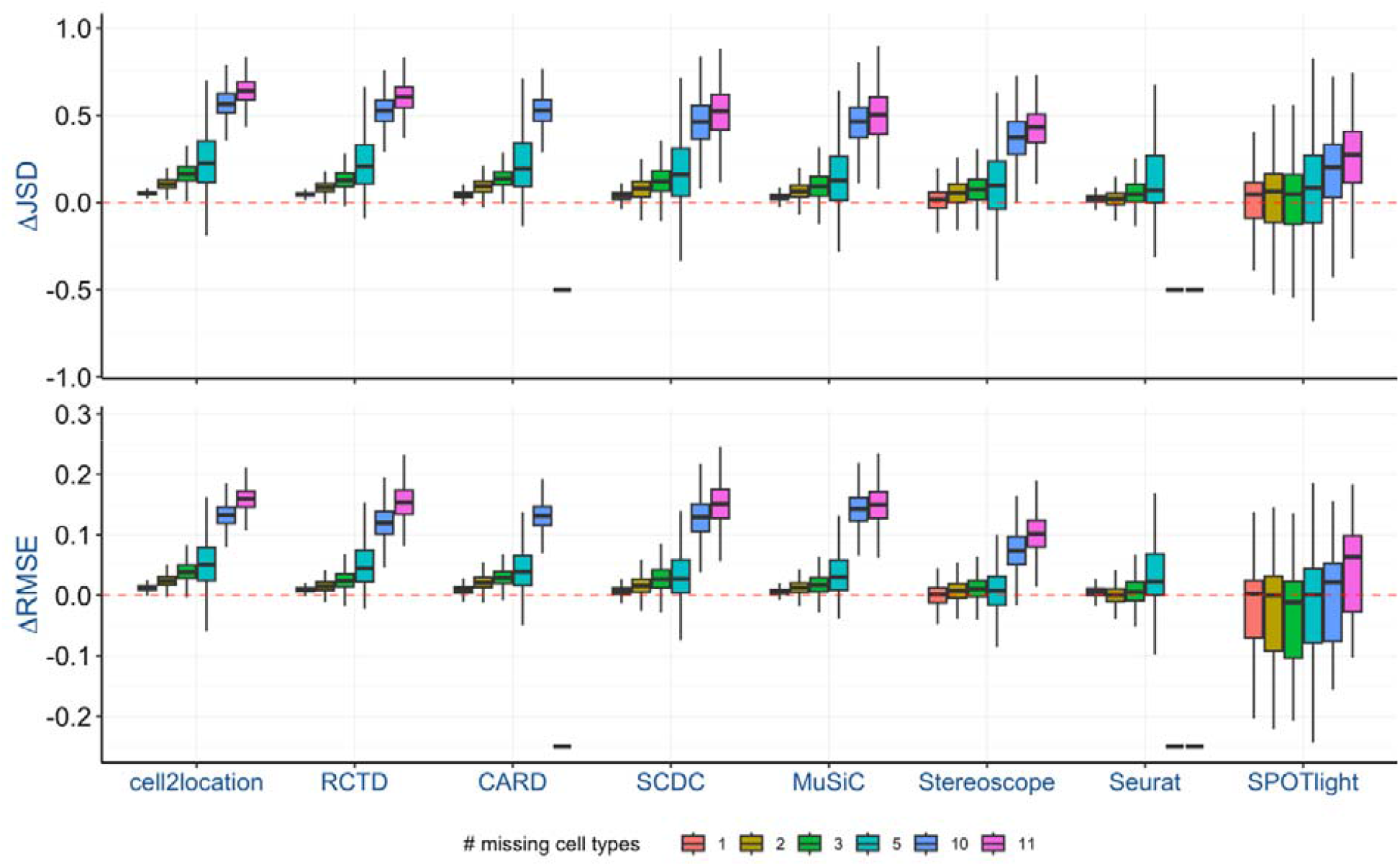
Performance of deconvolution methods in case of cell type mismatch for the lymph node data. Boxplots of the spot-wise difference in performance (ΔJSD, ΔRMSE) between the scenario where cell types are missing from the reference data and the baseline scenario for simulated dataset ST1. A value of ΔJSD/ΔRMSE (y-axis) above zero corresponds to a decrease in performance compared to the baseline scenario. The colour key indicates the number of missing cell types (see Supplementary Information S4, Table S2 . ΔJSD/ΔRMSE were calculated as the spot-wise mean across multiple instances of a particular scenario. A dash (-) indicates missing results for that particular scenario. Deconvolution methods are in the same order as in Figure 2. Deconvolution accuracy decreases as more cell types are removed from the reference data.

### 3.3 Proportions of missing cell types are assigned to transcriptionally similar cell types

The performance measures used until now do not differentiate between the cell types to which the proportions of the missing cell type(s) are assigned. We therefore calculated the reassignment value that captures to which cell type(s) the proportions of the removed cell type(s) are reassigned compared to the baseline scenario (see Supplementary Information S4). The reassignment values when leaving out one cell type from the reference data are visualized in Figure 4a. For the top-performing methods at baseline (cell2location, RCTD, CARD), similar trends can be identified across most removed cell types. We observe that (*i*) proportions for adipocytes are mostly reassigned to those for myocytes, (*ii*) proportions for CD4 T cells are mostly reassigned to those for CD8 T cells, (*iii*) proportions for CD8 T cells are more or less equally reassigned to CD4 T cells and NK cells, (*iv*) proportions for endothelial cells and fibroblasts are mostly reassigned to adipocytes, (*v*) proportions for macrophages are mostly reassigned to monocytes, (*vi*) proportions for myocytes are more or less equally reassigned to adipocytes and skeletal muscle cells, (*vii*) proportions for NK cells are mostly reassigned to CD8 T cells, and (*viii*) proportions for skeletal muscle cells are mostly reassigned to myocytes. Note that reassignment is not necessarily symmetric, as can be seen for adipocytes on the one hand and endothelial cells/fibroblasts on the other hand, for example. For the other cell types there is only partial agreement among top-performing methods. B cells are mostly reassigned to CD4 T cells by cell2location, whereas reassignment is spread out over various cell types for RCTD and CARD. Haematopoietic stem cells (HSCs) are mostly reassigned to NK cells by cell2location and CARD, but more or less equally reassigned to CD4 T cells and NK cells by RCTD. Monocytes are mostly reassigned to macrophages by RCTD and to B cells by CARD, whereas reassignment is spread out over various cell types (including B cells and macrophages) for cell2location. Neutrophils are mostly reassigned to monocytes by RCTD, whereas the reassignment is spread out over a large number of cell types for cell2location and CARD. Results for the other two simulated ST datasets showed similar trends (Figure S6).

**Figure 4.**
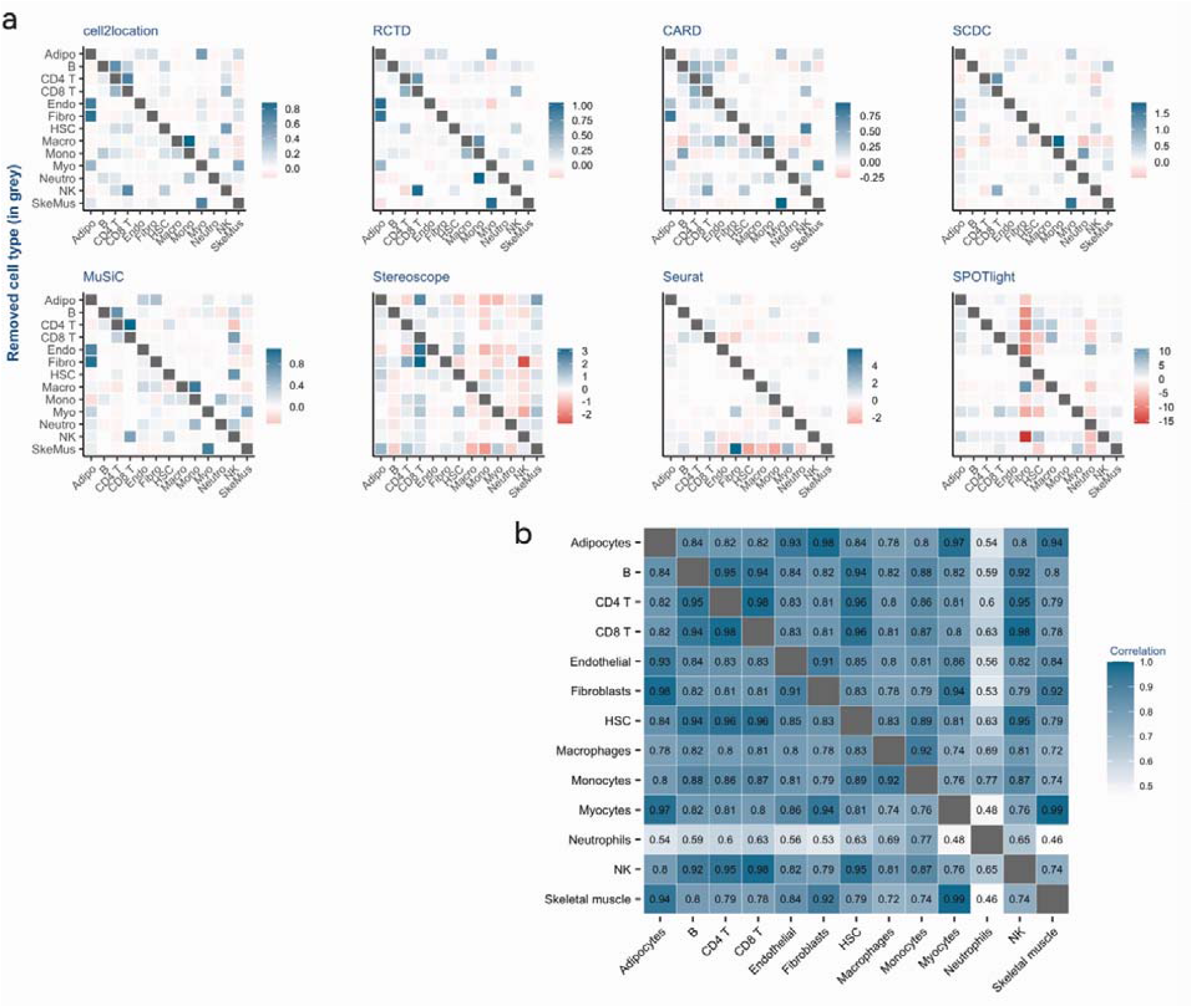
Cell-type-specific effects for the mismatch scenario with one cell type missing from the lymph node reference data for simulated dataset ST1. (a) Heatmaps of the reassignment values for each deconvolution method. For a cell type excluded from the reference data, the reassignment value denotes the normalized change in predicted proportions, relative to the baseline scenario proportions, for each of the remaining cell types (see Supplementary Information S4). Each row of the heatmap displays the reassignment values for the cell types indicated on the x-axis, following the removal of the cell type indicated on the *y*-axis (highlighted by the grey cell). Positive values represent an increase in proportion, negative values represent a decrease in proportion (as indicated by the colour key). Deconvolution methods are in the same order as in Figure 2. (b) Heatmap of Pearson correlation coefficients of the mean expression profiles of the indicated cell types using the 3,000 most highly variable genes. Adipo: adipocytes; Endo: endothelial cells; Fibro: fibroblasts; HSC: haematopoietic stem cells; Macro: macrophages; Mono: monocytes; Myo: myocytes; Neutro: neutrophils; SkeMus: skeletal muscle cells. For the better-performing methods, missing cell type proportions are mainly reassigned to transcriptionally similar remaining cell types.

As can be seen from the pairwise Pearson correlation between the mean expression profiles of the cell types in the reference data (Figure 4b), for the top-performing methods the cell types are in general reassigned to the transcriptionally most similar cell type(s). For example, CD4 T cells are most strongly correlated with CD8 T cells, with a correlation value of 0.98. CD8 T cells are most strongly correlated with CD4 T cells and NK cells with a correlation value of 0.98. Similarly, endothelial cells and fibroblasts are most strongly correlated with adipocytes with a correlation value of 0.93 and 0.98, respectively. Also, skeletal muscle cells are most strongly correlated with myocytes with a correlation value of 0.99. Some exceptions can also be observed. For example, adipocytes are most strongly correlated with fibroblasts with a correlation value of 0.98, followed by myocytes with a correlation value of 0.97. In this case, the reassignment values show a slightly different ranking with proportions for adipocytes mostly reassigned to those for myocytes and with fibroblasts being ranked lower, but still in the top 3.

The worst performing methods (Stereoscope, Seurat, SPOTlight) show a completely different picture. In this case, the proportion of cells originally assigned to the now missing cell type is often reassigned to multiple different cell types, and not necessarily to the most transcriptionally similar cell type. Moreover, it is often accompanied by a substantial decrease in the proportions of various cell types (Figure 4a, red squares). This corresponds to a widespread redistribution of the cell type proportions compared to the baseline scenario, even when removing just a single cell type from the reference data. A typical example is when leaving out skeletal muscle cells for Stereoscope. This leads to a marked increase in proportions for adipocytes, CD4 T cells, and myocytes. At the same time, proportions for macrophages and monocytes show a substantial decrease. Given the mediocre results for these methods in the baseline scenario, this is hardly surprising. Since the predicted proportions already substantially deviate from the ground truth in the baseline scenario, major shifts when leaving out one (or more) cell type(s) are to be expected.

We also extended our analysis to the removal of two (Figure S7) or three (Figure S8) cell types from the reference data. These show similar trends, with missing cell types assigned to the transcriptionally most similar cell type(s) for the top-performing methods and a major redistribution of cell type proportions for the worst performing methods. For example, when removing both adipocytes and fibroblasts, proportions are mostly reassigned to myocytes by cell2location, RCTD, and CARD (Figure S7). Myocyte expression is indeed strongly correlated with both adipocyte and fibroblast expression with a correlation value of 0.97 and 0.94, respectively (Figure 4b). Similarly, when removing NK cells, CD4 T cells, and CD8 T cells, proportions are mostly reassigned to HSCs (Figure S8), the expression of which is indeed strongly correlated with all three cell types.

### 3.4 Validation using a hypothalamus single-nucleus reference dataset

To assess whether the trends observed on the lymph node dataset generalize to another tissue/dataset, we performed all analyses on a hypothalamus single-nucleus dataset from a dissection of the supraoptic region [35]. Full details are provided in the Supplementary Information (Sections S5-S6, Tables S3-S4, Figures S10-S18).

In short, the results on the hypothalamus data largely confirm our main conclusions from the lymph node data. First, cell type deconvolution methods show large differences in baseline performance (Figure S11). Furthermore, the ranking of the deconvolution methods at baseline is largely preserved, with cell2location being the top-performing method and Stereoscope, Seurat, and SPOTlight performing substantially worse. Compared to the lymph node dataset, performance of MuSiC improved and performance of RCTD slightly deteriorated. Second, deconvolution performance decreases proportionally to the number of cell types missing from the reference data (Figure S13-S14). Third, proportions of missing cell types are assigned to transcriptionally similar cell types (Figure S15-S17). However, reassignment patterns on the hypothalamus data are slightly more diffuse across methods than on the lymph node data.

In summary, these results support the generalisability of the main conclusions of our study, while also indicating that the exact rankings of the deconvolution methods are partly dataset-dependent.

## 4. Discussion

Our study aimed to systematically evaluate and compare the robustness of different reference-based deconvolution methods for ST data, especially when the scRNA-seq reference used has one or more missing cell types. For this purpose, we included six methods specifically tailored for ST data and two methods commonly used for the deconvolution of bulk RNA-seq data. The combined results for simulated dataset ST1 generated from the lymph node and hypothalamus reference datasets are summarized in Figure 5, while corresponding dataset-specific summaries for lymph node (for simulated datasets ST1-3) and hypothalamus data (for simulated dataset ST1) are shown in Figure S9 and S18, respectively. Firstly, for the baseline scenario with no cell types missing from the reference, cell2location outperformed the other methods with RCTD, CARD, and MuSiC also showing good performance. This is in agreement with other recent benchmarking studies [27, 28]. However, the ranking of the deconvolution methods is partly dataset-dependent with RCTD showing better baseline performance on the lymph node data (Figure 2, Figure S3a,c) and MuSiC on the hypothalamus data (Figure S11). Secondly, performance decreased proportionally with the number of cell types missing from the reference. Therefore, cell2location showed the best overall performance both with and without missing cell types with RCTD, CARD, and MuSiC also performing relatively well. The performance of the other deconvolution methods was more variable, with SCDC showing average performance and SPOTlight consistently performing worst. Note that in practice deconvolution performance may not be the only criterion to select a specific method. Often the computational cost is also an important practical consideration. A quantitative comparison on ST1 derived from the lymph node reference data in the baseline scenario shows substantial differences in run time for the top-performing methods with CARD (∼1 minute) and RCTD (∼3 minutes) being fast and cell2location (∼17 minutes) and especially MuSiC (∼38 minutes) requiring substantially more time (Table S5). Note that run time for cell2location is on GPU, since of these four methods it is the only one that allows for GPU acceleration. Therefore, we conclude that across a range of scenarios cell2location is the most effective method if accuracy and robustness are most important and GPU is available, otherwise, if run time on CPU is also an important consideration, RCTD and CARD are good options.

**Figure 5.**
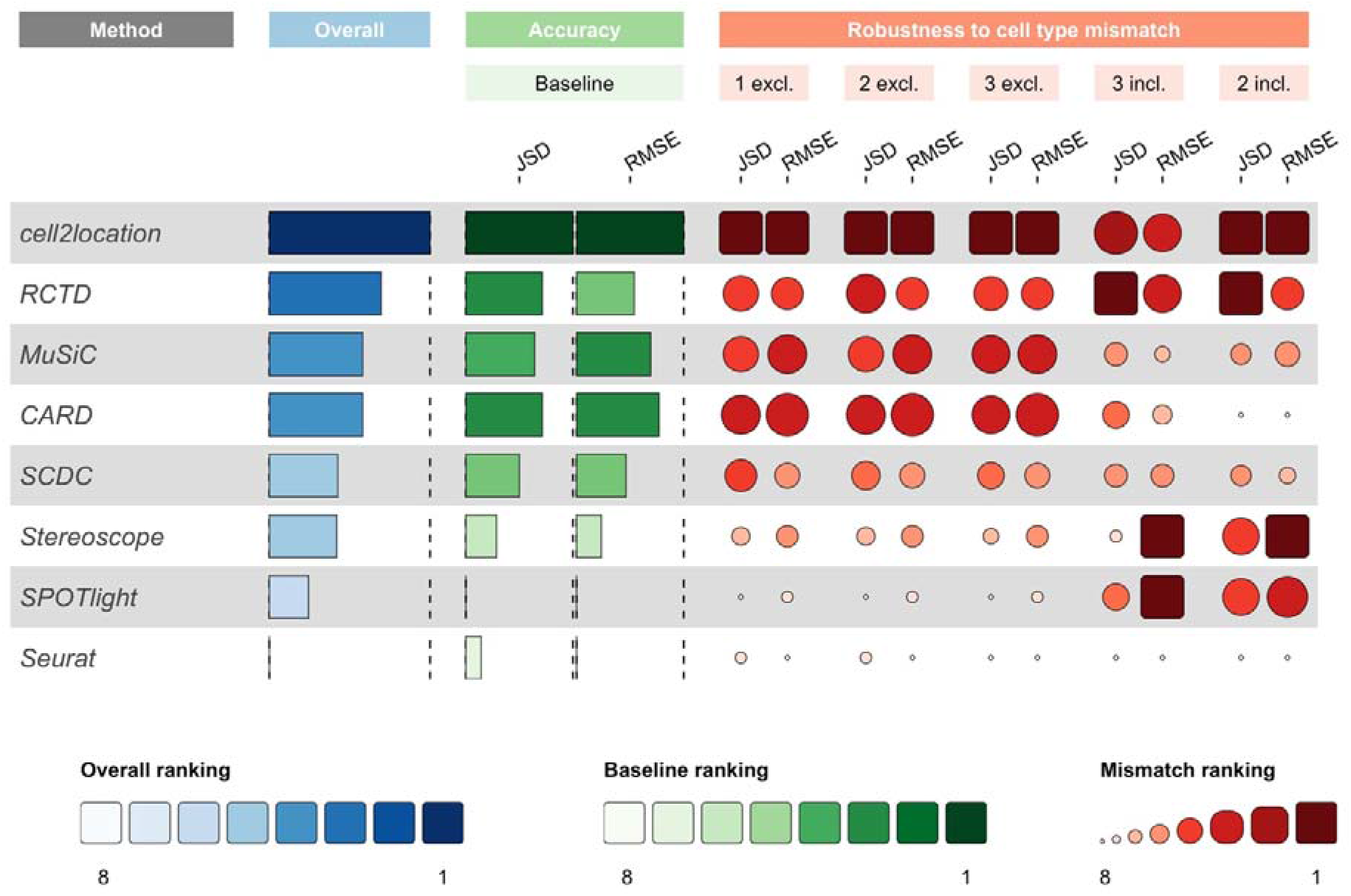
Summary of deconvolution results for simulated dataset ST1 generated from the lymph node and hypothalamus reference datasets. Performance is shown for the baseline scenario (accuracy; green) and in case of cell type mismatch (robustness to cell type mismatch; red) and summarized as overall performance (blue). Deconvolution methods are ranked using the mean value of the indicated performance metrics (JSD, RMSE). Darker shades indicate better performance. The overall ranking was computed using the mean of all baseline and mismatch scenario rankings for both reference datasets (for both JSD and RMSE). For the mismatch scenarios, we only included those shared between the lymph node and hypothalamus dataset. ‘excl.’ indicates the number of cell types missing from the reference dataset, ‘incl.’ indicates the number of cell types remaining in the reference dataset. The overall trends are consistent across datasets and show that cell2location is the most effective method in terms of accuracy and robustness.

We also investigated the consequences of missing cell types on the biological interpretation of the results. For this purpose, we quantified how predicted cell proportions are redistributed using the reassignment value. Across a range of scenarios with 1-3 cell types missing from the reference datasets, this clearly showed that proportions of missing cell types are assigned to transcriptionally similar cell types. This is in line with earlier observations for a highly specific example with one or two cell types missing from the reference data for RCTD [20]. Redistribution of cell type proportions has been studied in more detail in the context of deconvolution of bulk RNA-seq data [36, 37]. There, it was also shown that missing cell type proportions were assigned to the transcriptionally most similar cell type(s). However, these studies only investigated the scenario with one cell type missing from the reference data. Recently, Ivich et al. performed an in-depth study of the influence of cell types from a single-cell reference on the deconvolution of bulk RNA-seq data [38]. They observed that the profiles of the missing cell types can to a certain extent be recovered from residuals using non-negative matrix factorization. Whether similar observations can be made in the context of deconvolution of ST data remains to be investigated.

Our results are based on data that we simulated using a large lymph node scRNA-seq reference dataset [32, 33] and a hypothalamus snRNA-seq reference dataset [35]. For this purpose, we developed an algorithm for simulating ST data that enabled us to investigate various scenarios by varying the total number of cells and cell types present per spot, as well as the sequencing depth. In our simulation approach, each spatial location is considered independently without taking into account the possible spatial correlation between neighbouring spots typically observed in a profiled tissue. Approaches that simulate ST data corresponding to pre-specified synthetic tissue patterns are starting to be developed [39]. However, note that of the eight deconvolution methods that we evaluated only CARD can explicitly exploit that nearby spots often contain similar cell types, using a conditional autoregressive model [10]. Our simulation approach therefore aligns with our primary objective, which is to create a reliable starting point that focuses solely on identifying the cell types present in each spot using the reference data. As future work, the algorithm can be expanded to incorporate spatial correlations between spots. At the same time, this means that our systematic evaluation was performed in a setting where CARD’s spatial modelling component confers no advantage and its performance in real tissues with spatial structure may therefore be underestimated.

Our results clearly show that current deconvolution methods for ST data lack robustness in the commonly occurring scenario where cell types are missing from the single-cell reference data. There are several approaches to increase robustness to cell type mismatch. It has been suggested to crop the ST data and exclude regions with cell types that are only present in the ST data and not in the scRNA-seq reference data [20]. Alternatively, a recent study proposed utilizing sorted scRNA-seq data to create a cell mixture resembling the ST data and use that to predict the spatial cell composition when cell types are missing from the reference data [30]. However, these approaches still assume prior knowledge of the structure and cell type composition of the profiled tissue. While reference-based deconvolution methods cannot identify entirely new cell types, future approaches could enhance robustness by explicitly modelling an unknown component and estimating uncertainty in the predicted proportions. At the same time, it is important to note that current deconvolution tools have limited ability to accurately resolve ST data in case of cell type mismatch, and researchers should interpret such results with appropriate caution. Therefore, we advocate for the development of generic reference-based cell type deconvolution methods that are more robust to cell type mismatches. Our framework is implemented in a modular way and can readily be extended to evaluate such newly developed deconvolution methods.

## Key Points

– Cell2location outperforms other methods, revealing clear differences in baseline deconvolution accuracy.
– Performance consistently worsens as more cell types are missing from the reference, revealing a fundamental vulnerability in current deconvolution strategies.
– Missing cell types are reassigned to transcriptionally similar cell types in top-performing methods, highlighting predictable behaviour but also the need for more robust approaches.

## Data availability

All data used in this study are publicly available. The Tabula Sapiens lymph node single-cell dataset can be downloaded from https://figshare.com/articles/dataset/Tabula_Sapiens_release_1_0/14267219/3 (TS_Lymph_Node.h5ad.zip; version 3). The lymph node stromal single-cell data are available at GEO with accession number GSE261747 (https://www.ncbi.nlm.nih.gov/geo/query/acc.cgi?acc=GSE261747).

The human hypothalamus single-cell dataset used for the independent validation was downloaded from the Human Single-Cell Atlas (https://data.humancellatlas.org/hca-bio-networks/nervous-system/atlases/brain-v1-0).

## Code availability

The code to reproduce all analyses and results is available at https://github.com/EDS-Bioinformatics-Laboratory/Robustness_evaluation_deconv_methods.

## Acknowledgements

We thank Cristoforo Grasso for providing early access to the lymph node stromal cell single-cell dataset. We thank all members of the Bioinformatics Laboratory at the Amsterdam UMC for helpful discussions. This study was supported by ARCAID (www.arcaid-h2020.eu) which has received funding from the European Union’s Horizon 2020 research and innovation program under the Marie Skłodowska-Curie grant agreement no 847551. For computational analyses this study made use of the Dutch national e-infrastructure with the support of the SURF Cooperative using grant no. EINF-3461.

