## Supplementary Information for "Systematic evaluation of robustness of deconvolution methods for spatial transcriptomics data in case of cell type mismatch"

^2^ Amsterdam Public Health, Methodology, Amsterdam, The Netherlands

^3^ Amsterdam institute for Immunology & Infectious Diseases, Inflammatory Diseases, Amsterdam, The Netherlands

^4^ Amsterdam UMC location University of Amsterdam, Rheumatology and Clinical Immunology, Meibergdreef 9, Amsterdam, The Netherlands

^5^ Amsterdam Rheumatology & Immunology Center (ARC), Amsterdam, The Netherlands

*Corresponding author:

Perry D. Moerland

Bioinformatics Laboratory

Epidemiology and Data Science

Amsterdam University Medical Centers

Meibergdreef 9, 1105 AZ Amsterdam, The Netherlands

**Supplementary Information**

**S1. Lymph node single-cell reference data**

We used two human lymph node scRNA-seq datasets to construct an integrated reference dataset. The Tabula Sapiens Consortium lymph node data (version 3) [1] comprises 53,275 cells, most of which are B or T cells. Since this dataset contains only few fibroblasts, we also included a lymph node stromal cell (LNSC) dataset consisting of 13,850 cells [2]. The datasets were further processed and integrated using the Seurat (v4.1.0) [3] R package with R version 4.1.2.

**S1.1 Processing of Tabula Sapiens lymph node data.** We excluded cells measured using the Smart-seq2 protocol and selected the remaining 50,981 cells measured using 10x Genomics platforms. We reannotated the remaining cells with the SingleR R package (v1.10.0) [4] using normalized expression values of bulk RNA-seq samples generated by Blueprint and ENCODE from pure populations of stroma and immune cells [5, 6] as reference data (function ‘BlueprintEncodeData’; R package celldex (version 1.4.0)). Annotation was done at the level of 24 broad classes (‘label.main’) and using the pruned labels in order to exclude low-quality annotations. The original Tabula Sapiens uniform manifold approximation and projection (UMAP) coordinates were used to visualize the data for all three donors combined (Figure S1a). Next, the data were split into four batches based on the donor and 10x platform (3’ or 5’) from which the cells originated. Further processing steps were performed separately for each batch. To discard low-quality cells we selected cells with at least 750 detected genes (nFeature_RNA), at least 2,300 counts (nCount_RNA), a novelty score (log10GenesPerUMI) greater than 0.8 [7], and less than 30% mitochondrial counts. The number of cells after filtering was 49,405 (donor 1: 8,351, donor 2: 11,503, donor 3: 29,551 cells). Subsequently, the count data were log-normalized with a scaling factor of 10,000 and the 3,000 most highly variable genes were identified using the ‘vst’ method and standardized the expression per gene (function ‘ScaleData’).

**S1.2 Processing of lymph node stromal cell data.** To discard low-quality cells, we selected cells with at least 250 detected genes (nFeature_RNA), at least 500 counts (nCount_RNA), a novelty score (log10GenesPerUMI) greater than 0.7, and less than 20% mitochondrial counts. The number of cells after filtering was 12,298. Subsequently, the count data were log-normalized with a scaling factor of 10,000. We identified the 3,000 most highly variable genes using the ‘vst’ method and standardized the expression per gene (function ‘ScaleData’). Principal component analysis (PCA) was performed on the scaled data using 30 principal components (PCs). The PCs were used to determine the UMAP coordinates, which were used to visualize the data (Figure S1b). Cell annotation was performed as described above for the Tabula Sapiens data.

**S1.3 Data integration.** To combine the resulting five datasets we used Seurat’s anchor-based integration strategy based on canonical correlation analysis and a mutual nearest neighbour search [8]. First we selected the 3,000 most highly variable genes across the five datasets (function ‘SelectIntegrationFeatures’). Next, we identified integration anchors using these selected genes (function ‘FindIntegrationAnchors’) and then integrated the datasets (function ‘IntegrateData’). The resulting integrated data is comprised of 61,703 cells. We then performed scaling and PCA on the scaled data using 30 PCs. See Figure S1c for a UMAP plot of the integrated data.

**S1.4 Downsampling of cells.** In order to reduce the computation time required for benchmarking the deconvolution methods, we selected cells from the integrated data. First, we removed cells for which the annotation label was missing. Since the cell type distribution was highly non-uniform (Table S1), we selected the 13 cell types with at least 95 annotated cells. Next, cell types with more than 300 annotated cells were randomly downsampled to 300 cells (Table S1). The resulting dataset consists of 3,519 cells distributed across 13 cell types (Figure S1d).

**S1.5 Gene selection.** To further reduce computation time we also adopted the gene filtering strategy proposed by Kleshchevnikov et al. [9]. We selected genes expressed (count ≥ 1) in at least 3% of the cells and, in addition, genes expressed in 0.01% to 3% of the cells but with a mean expression across cells expressing the genes greater than 2. We refer to the resulting dataset as the reference dataset, which comprises 11,474 genes and 3,519 cells.

**S2. Cell type deconvolution methods**

In our analysis, we included six cell type deconvolution methods tailored for ST data as well as two methods tailored for bulk RNA-seq data. Here, we briefly summarize the methodology of each method and choices we made for their application and parameter values.

**CARD** (v1.0) [10]: This method uses non-negative matrix factorization (NMF) for deconvolution of ST data given cell type-specific gene expression in a scRNA-seq reference dataset. CARD exploits that nearby spots often contain similar cell types using a conditional autoregressive model. Thus, we also provided the spatial coordinates of the simulated ST data as input. Note, however, that our simulated ST data has no spatial correlation structure and that, therefore, CARD is evaluated in what the developers refer to as a high-noise scenario [10]. For our analysis, we followed the guidelines provided by the developers of CARD (date accessed: 06/09/2022): <https://github.com/YMa-lab/CARD/blob/master/docs/documentation/04_CARD_Example.md>. Even though our single-cell reference data comprises cells from five batches, we considered it to be a single sample and indicated this in the associated metadata. All other parameters were set to the values suggested in the tutorial.

**cell2location** (v0.1) [9]: This method uses a hierarchical Bayesian model to estimate the abundance of each cell type at each spot by decomposing the spatial expression count matrix using cell type signatures estimated from the scRNA-seq reference dataset. For the analysis, we followed the scvi-tools [11] guidelines for the cell2location model (date accessed: 06/09/2022):

<https://docs.scvi-tools.org/en/0.16.2/tutorials/notebooks/cell2location_lymph_node_spatial_tutorial.html>. First, reference cell type signatures were estimated with a negative binomial regression model setting max_epochs = 1000. Next, the spatial mapping model for spatial data was trained with max_epochs = 10000 and hyperparameter detection_alpha = 200. The hyperparameter ‘N_cells_per_location’ was set based on the average number of cells present per spot in our simulated datasets (see Section S4), that is either equal to 13 (ST1 and ST2) or 5 (ST3). We used the 5% quantile of the posterior distribution of the cell type abundances per spot and then normalized these values so that their spot-wise sum is equal to 1.

**RCTD** (spacexr, v2.0.0) [12]: This method infers cell type proportions using maximum likelihood estimation assuming that gene counts are Poisson distributed. We followed the guidelines for applying RCTD to ST data provided by the method developers (date accessed: 06/09/2022):

[https:/‌/‌raw.githack.com/‌dmcable/‌spacexr/‌master/‌vignettes/‌spatial-transcriptomics.html](https://raw.githack.com/dmcable/spacexr/master/vignettes/spatial-transcriptomics.html). RCTD was used in full mode (i.e., doublet_mode = ‘full’) where there is no restriction on the number of cell types in a spot.

**Seurat** (v4.1.0) [8]: We used Seurat's approach to annotate the ST (query) dataset based on the cell type labels of the scRNA-seq (reference) dataset following the guidelines provided by the method developers (date accessed: 06/09/2022): <https://satijalab.org/seurat/archive/v3.2/integration#standard-workflow> [3] In short, we first log-normalized the scRNA-seq and ST data with a scaling factor of 10,000 and identified the 3,000 most highly variable genes in the reference data using the ‘vst’ method. Next, anchors between the scRNA-seq and ST datasets were determined in the canonical correlation embedding of both datasets (function 'FindTransferAnchors'). The set of calculated anchors was then used to transfer cell type annotation from the reference data to the ST dataset (function 'TransferData'). The resulting prediction scores were normalized so that their spot-wise sum is equal to 1.

**SPOTlight** (v1.0.0) [13]: This method uses a combination of NMF regression and non-negative least squares to deconvolve ST data. We followed the guidelines provided in the vignette by the method developers (date accessed: 06/09/2022): [https:/‌/‌github.com/‌MarcElosua/‌SPOTlight/‌blob/‌main/‌vignettes/‌SPOTlight_kidney.Rmd](https://github.com/MarcElosua/SPOTlight/blob/main/vignettes/SPOTlight_kidney.Rmd).

No additional cell downsampling was performed and a marker gene score cut-off of 0.5 (mean.AUC > 0.5) was used.

**Stereoscope** (scvi-tools v0.16.2) [14]: This method uses a negative binomial distribution to model scRNA-seq and spatial data. We used the implementation of Stereoscope available in scvi-tools [11] following their guidelines (date accessed: 06/09/2022):

<https://docs.scvi-tools.org/en/0.16.2/tutorials/notebooks/stereoscope_heart_LV_tutorial.html>. We did not perform any additional filtering steps for both datasets. All other parameters were set to the values suggested in the tutorial.

**MuSiC** (v1.0.0) [15]: This method has been developed for deconvolution of bulk RNA-seq data with scRNA-seq reference data using weighted non-negative least squares (W-NNLS) regression. We followed the guidelines provided by the method developers for single sample data (date accessed: 06/09/2022): [https:/‌/‌xuranw.github.io/‌MuSiC/‌articles/‌MuSiC.html](https://xuranw.github.io/MuSiC/articles/MuSiC.html). Cell type proportions were estimated without pre-grouping of cell types and no prior selection of marker genes was performed. We considered each spot in the ST data as one bulk RNA-seq sample, resulting in 1600 samples.

**SCDC** (v0.0.0.9000) [16]: This method has also been developed for the deconvolution of bulk RNA-seq data with scRNA-seq reference data using w-NNLS. We followed the guidelines provided by the method developers (date accessed: 06/09/2022): [https:/‌/‌meichendong.github.io/‌SCDC/‌articles/‌SCDC.html](https://meichendong.github.io/SCDC/articles/SCDC.html). In specific, we followed the three-cell-line mixture data analysis example for single sample data using the function `SCDC_prop_ONE`. We considered each spot in the ST data as one bulk RNA-seq sample, resulting in 1600 samples. All parameters were used with default values except the ‘weight.basis’ parameter, which was set to ‘FALSE’ to avoid SCDC generating a basis matrix with NA values.

**S3. Generation of simulated spatial transcriptomics data**

To benchmark the deconvolution methods outlined above, we developed a procedure to generate ST data with known per spot cell type proportions from a given single-cell reference dataset. Our simulation procedure generates ST data with 1,600 spots where both the number of cells and the number of cell types per spot can be varied to assess the effect of different spot sizes and different heterogeneity of the underlying tissue.

For a reference dataset consisting of *C* cell types, we first draw the number of cell types *C_i_* in spot *i* from a discrete uniform distribution *U*{*C*_min_, *C*_max_} and then randomly sample a set *S_i_* of *C_i_* cell types (Figure S2). Next, we determine the proportion of each cell type present in a spot by *C_i_* independent draws from a continuous uniform distribution *U*(0,1). The resulting values are normalized such that their sum equals 1, giving proportions *P_i,k_* of cell type *k*. Subsequently, the number of cells *D_i_* in a spot is drawn from a discrete uniform distribution *U*{*D*_min_, *D*_max_}. Next, using both the cell type proportions *P_i,k_* and the number of cells *D_i_*, we determine the number of cells per cell type *W_i,k_* by sampling from a multinomial distribution *M*{*D_i_; P_i,1,…,_ P_i,C_*}. The corresponding proportions *W_i,k_*/*D_i_* serve as the ground truth for our benchmarking study. We then simulate the mRNA counts *M_i,g_* for gene *g* using the following equation:

$$M_{i,g}= \sum_{k=1}^{C_{i}} R_{g,k}* W_{i, k}$$

Here, *R_g,k_* denotes the mRNA count for gene *g* in cell type *k* (in spot *i*). *R_g,k_* is determined per spot by calculating the ceiling of the average of the counts for gene *g* in five cells of cell type *k* randomly drawn from the reference dataset. Finally, to make the total number of counts per spot similar to what is observed in 10x Visium ST data, for those spots with more than 30,000 counts we draw a random number from a discrete uniform distribution *U*{25000, 30000} and scale the per gene counts accordingly (function ‘downsampleMatrix’, R package scuttle). We also generate spatial coordinates for each spot in a 40 by 40 layout for those deconvolution methods that require the spot coordinates as input.

**S4. Benchmark framework and performance evaluation**

To systematically evaluate the robustness to cell type mismatch of the eight deconvolution methods described in Section S2, we designed a benchmark framework based on simulated ST data. The integrated human lymph node data (Section S1) were used both as single-cell reference dataset and as input for simulating ST data with known per spot cell type proportions. With the simulation approach described in Section S3 we generated three ST datasets from the lymph node reference data varying both the number of cell types (C) and the number of cells (D) present per spot with the following values for (*C*_min_, *C*_max_) and (*D*_min_, *D*_max_):

- ST1: 4-8 cell types and 10-15 cells per spot;
- ST2: 1-5 cell types and 10-15 cells per spot;
- ST3: 1-5 cell types and 3-7 cells per spot, with the number of cells greater than or equal to the number of cell types.

The proportions estimated by each deconvolution method were then compared with the simulated per spot cell type proportions *W_i,k_*/*D_i_* (that is, the ground truth). First, we evaluated deconvolution performance for the baseline scenario without cell type mismatch, that is, both the single-cell reference and the simulated ST data contain the same cell types. Next, we assessed robustness to cell type mismatch by systematically removing one or more cell types from the reference data, but not from the ST data. We investigated the following cell type mismatch scenarios indicated by the number of removed cell types (Table S2):

- 1 type: Sequential removal of each of the 13 cell types (13 simulations);
- 2 types: Sequential removal of five pairs of transcriptionally similar cell types (5 simulations);
- 3 types: Sequential removal of five triplets of transcriptionally similar cell types (5 simulations);
- 5 types: removal of one quintuplet (1 simulation);
- 10 types: complementary to scenario 3, sequential removal of five sets of ten cell types not included in the triplet (5 simulations);
- 11 types: complementary to scenario 2, sequential removal of five sets of eleven cell types not included in the pairs (5 simulations).

Both for the baseline and cell type mismatch scenarios we quantified the performance of the each of the deconvolution methods by comparing the ground truth per spot cell type proportions and the predicted proportions. For this purpose we selected two commonly used performance measures: Jensen-Shannon divergence (JSD) and root mean square error (RMSE). In addition we also define the reassignment value that captures to which cell type(s) the proportions of the removed cell type(s) get reassigned.

**Jensen-Shannon divergence**: The JSD measures the similarity between two probability distributions (here, ground truth (*gt*) and predicted proportions (*pred*) in a spot). The JSD is a symmetrized version of the Kullback-Leibler divergence (KLD). We used the following equations to calculate the JSD for each spot:

$$\mathrm{KLD}\left( P | | Q \right)= \sum_{k=1}^{C} P_{k} \log_{2} \left( \frac{P_{k}}{Q_{k}} \right)$$

$$M= \frac{1}{2} \left( pred+gt \right)$$

$$\mathrm{JSD}\left( pred | | gt \right)= \frac{1}{2} \mathrm{KLD}\left( pred | | M \right)+ \frac{1}{2} \mathrm{KLD}\left( gt | | M \right)$$

Here vectors *pred* and *gt* denote the predicted and ground truth cell type proportions of *C* cell types, respectively. The JSD is bounded between 0 and 1, and equal to zero if and only if the two distributions are identical. Therefore, a lower JSD indicates better performance.

**Root mean square error**: We used the following equation to calculate the RMSE for each spot:

$$\mathrm{RMSE}= \sqrt{\frac{\sum_{k=1}^{C} {{(pred}_{k}- {gt}_{k})}^{2}}{C}}$$

A lower RMSE indicates better performance.

**Reassignment value:** The predicted proportions of a cell type at baseline become zero if the cell type is removed from the reference data in one of the cell type mismatch scenarios. Consequently, this is accompanied by an increase of the predicted proportions of one or more of the remaining cell types. Therefore, we define the reassignment value to capture to which cell type(s) the proportions of the removed cell type(s) are reassigned. The calculation of the reassignment value is outlined in Algorithm S1.

**Algorithm S1: Calculation of reassignment values**

**B**: 1600 x 13 matrix of predicted cell type proportions at baseline for 1600 spots and 13 cell types

**P**: 1600 x (13 – *j*) matrix of predicted cell type proportions with $j\in$ {1,2,3,5,10,11} cell types removed from the reference data

*S*: set of remaining cell types, that is |*S*| = 13 – *j*

*R*: set of removed cell types, that is |*R*| = *j*

**for** *k* **in** *S*

\# Calculate change in proportions w.r.t. the baseline scenario for cell type *k* for all spots

***d*** = **P**[:, *k*] - **B**[:, *k*]

\# Calculate the reassignment value *rv* for cell type *k* when removing the cell types in *R*

\# Here, the numerator corresponds to the total change in proportions of cell type *k* w.r.t. to baseline

\# and the denominator corresponds to the sum of the proportions at baseline of the removed cell types

rv*_R,k_* = $\sum_{i=1}^{1600} d_{i} /\sum_{i=1}^{1600} \sum_{r \in R} \mathbf{B}[i,r]$

**end**

The reassignment value rv*_R,k_* corresponds to the change in predicted proportions w.r.t. the baseline scenario for a given cell type *k* when removing the cell types contained in *R* from the reference data, normalized by the sum of the proportions at baseline of the removed cell types. This means that if the proportions of the removed cell types *R* are all reassigned to cell type *k,* its reassignment value rv*_R,k_* is equal to one and the reassignment values of the other remaining cell types are equal to zero. Note, however, that in practice reassignment values can be larger than one or even negative, since the removal of a set of cell types will in general affect the predicted proportions for all remaining cell types.

**Examples of reassignment value calculation**: We consider a minimal example with three spots and three cell types (A, B, C), where we remove cell type C (so, *R* = {*C*} and *S* = {*A, B*}). Assume that the matrix of predicted proportions at baseline is:

| **Spot** | **A** | **B** | **C** |
| --- | --- | --- | --- |
| 1 | 0.2 | 0.3 | 0.5 |
| 2 | 0.1 | 0.4 | 0.5 |
| 3 | 0.3 | 0.2 | 0.5 |

The total proportion at baseline of cell type C over all three spots is 1.5, corresponding to the denominator of the reassignment values. Now we will give examples of four scenarios when cell type C is removed.

Scenario 1: the reassignment value of A, rv*_C,A_*, is equal to one. This occurs when the total proportion of C is reassigned to A. Assume that the matrix of predicted proportions after removal of cell type C is:

| **Spot** | **A** | **B** |
| --- | --- | --- |
| 1 | 0.7 | 0.3 |
| 2 | 0.6 | 0.4 |
| 3 | 0.8 | 0.2 |

Then ${rv}_{C,A}=\frac{\left( 0.7-0.2 \right)+\left( 0.6-0.1 \right)+(0.8-0.3)}{1.5}=\frac{1.5}{1.5}=1$ and ${rv}_{C,B}=\frac{\left( 0.3-0.3 \right)+\left( 0.4-0.4 \right)+\left( 0.2-0.2 \right)}{1.5}=0$

Scenario 2: the reassignment value of A, rv*_C,A_*, is strictly between zero and one. This occurs when the total proportion of C is distributed across both A and B. Assume that the matrix of predicted proportions after removal of cell type C is:

| **Spot** | **A** | **B** |
| --- | --- | --- |
| 1 | 0.5 | 0.5 |
| 2 | 0.4 | 0.6 |
| 3 | 0.6 | 0.4 |

Then ${rv}_{C,A}=\frac{\left( 0.5-0.2 \right)+\left( 0.4-0.1 \right)+(0.6-0.3)}{1.5}=\frac{0.9}{1.5}=0.6$ and ${rv}_{C,B}=\frac{\left( 0.5-0.3 \right)+\left( 0.6-0.4 \right)+(0.4-0.2)}{1.5}= \frac{0.6}{1.5}= 0.4$

Scenario 3: the reassignment value of A, rv*_C,A_*, is greater than one. This occurs when the total proportion of A increases by more than the total proportion of C and is therefore accompanied by a decrease of the total proportion of B. Assume that the matrix of predicted proportions after removal of cell type C is:

| **Spot** | **A** | **B** |
| --- | --- | --- |
| 1 | 0.9 | 0.1 |
| 2 | 0.8 | 0.2 |
| 3 | 1.0 | 0.0 |

Then ${rv}_{C,A}=\frac{\left( 0.9-0.2 \right)+\left( 0.8-0.1 \right)+(1.0-0.3)}{1.5}=\frac{2.1}{1.5}=1.4$ and ${rv}_{C,B}=\frac{\left( 0.1-0.3 \right)+\left( 0.2-0.4 \right)+(0.0-0.2)}{1.5}= \frac{-0.6}{1.5}=-0.4$

Scenario 4: the reassignment value of A, rv*_C,A_*, is less than zero. This occurs when the total proportion of A decreases and is therefore accompanied by an increase of the total proportion of B by more than the total proportion of C. Assume that the matrix of predicted proportions after removal of cell type C is:

| **Spot** | **A** | **B** |
| --- | --- | --- |
| 1 | 0.1 | 0.9 |
| 2 | 0.0 | 1.0 |
| 3 | 0.2 | 0.8 |

Then ${rv}_{C,A}=\frac{\left( 0.1-0.2 \right)+\left( 0.0-0.1 \right)+(0.2-0.3)}{1.5}=\frac{-.0.3}{1.5}= -0.2$ and ${rv}_{C,B}==\frac{\left( 0.9-0.3 \right)+\left( 1.0-0.4 \right)+(0.8-0.2)}{1.5}= \frac{1.8}{1.5}=1.2$

**S5. Hypothalamus single-nucleus reference data**

As an independent validation dataset, we used a human hypothalamus single-nucleus RNA-seq dataset from a dissection of the supraoptic region [17], downloaded from the Human Single-Cell Atlas. The dataset comprised 12,557 neuronal and non-neuronal cells and 58,232 features and was converted from h5ad format into a Seurat object while preserving cell-level metadata (Figure S10).

Processing followed the same overall strategy as for the lymph node reference data (Section S1). Briefly, we used the provided supercluster annotations as cell type labels, retained cell types with at least 250 cells, and randomly downsampled cell types with more than 300 cells to 300 cells. This resulted in a reference dataset comprising 14 cell types (Table S3). Applying the same gene filtering strategy as before (Section S1.5) yielded a filtered dataset with 4,154 cells and 16,404 features (Figure S10), which was used both as the single-cell reference for deconvolution and as input for simulating ST data.

ST data were simulated using the framework described in Section S4, but only the simulated dataset corresponding to ST1 was generated. The same eight reference-based deconvolution methods were evaluated using the same workflows and parameter settings as in the main analysis.

As for the lymph node dataset, we considered the baseline scenario and multiple cell type mismatch scenarios in which one or more cell types were removed from the reference while keeping the ST data unchanged. Because the hypothalamus dataset had a different cell type composition, the mismatch scenarios were defined based on related hypothalamic populations. Specifically, we evaluated sequential removal of each of the 14 cell types, removal of selected related pairs, removal of selected related triplets, and complementary extreme mismatch scenarios in which only the selected triplets or only the selected pairs were retained in the reference (Table S4). Performance was evaluated as in the main benchmark using Jensen-Shannon divergence (JSD) and root mean square error (RMSE), together with changes relative to baseline and reassignment analysis (Section S4).

**S6. Supplementary results for the hypothalamus dataset**

**S6.1 Cell type deconvolution methods show large differences in baseline performance**

We first assessed the baseline scenario, in which both the hypothalamus single-nucleus reference and the simulated ST data contained the same 14 cell types. As for the lymph node data, clear performance differences between deconvolution methods were observed (Figure S11). Cell2location showed the best overall baseline performance, with the lowest median JSD (0.07) and median RMSE (0.04). MuSiC ranked second by median JSD (0.08) and first by median RMSE with similar value as that of cell2location (0.04). CARD, SCDC, and RCTD formed an intermediate-performing group, with median JSD values of 0.12, 0.12, and 0.13, respectively, and median RMSE values of 0.05, 0.06, and 0.07, respectively. In contrast, Stereoscope, Seurat, and SPOTlight performed substantially worse, with median JSD values of 0.37, 0.45, and 0.52, respectively, and median RMSE values of 0.09, 0.17, and 0.13, respectively. Thus, while on the lymph node data cell2location, RCTD, and CARD performed best, the hypothalamus data showed stronger relative performance of MuSiC and worse performance of RCTD.

To further characterize baseline performance, we also evaluated per-cell-type RMSE across the 14 hypothalamic cell types (Figure S12). The per-cell-type results were consistent with the method-level trends. Cell2location and MuSiC generally showed the lowest error across most cell types, whereas Seurat, SPOTlight, and Stereoscope showed a higher error across most cell types. Some cell types, such as committed oligodendrocyte precursor, appeared more challenging across methods than others. These results indicate that both overall method performance and cell-type-specific difficulty contribute to deconvolution accuracy in this dataset.

Overall, these results indicate that the relative ranking of methods at baseline is partly dataset-dependent, although the separation between better- and worse performing methods remained clear.

**S6.2 Performance decreases proportionally to the number of cell types missing from the reference data**

To evaluate robustness to cell type mismatch, we systematically removed one or more cell types from the hypothalamus reference data while keeping the simulated ST data unchanged, and subsequently applied the selected deconvolution methods. We considered cell type mismatch scenarios with 1, 2, 3, 11, or 12 missing cell types out of 14 cell types in total. In the case of one missing cell type, all 14 possible single-cell-type removals were evaluated. In the case of 2 or 3 missing cell types, we evaluated a limited number of combinations by defining sets of transcriptionally related hypothalamic cell types to be removed from the reference data (Table S4). We also included two extreme scenarios with 11 or 12 missing cell types, in which only 3 or 2 cell types remained in the reference, respectively. In these cases, only the selected triplets or pairs of transcriptionally related cell types were retained in the reference data. For all scenarios, spot-wise JSD and RMSE were calculated, and these were used to derive the spot-wise differences in performance relative to the baseline scenario (ΔJSD, ΔRMSE).

Both ΔJSD and ΔRMSE showed a clear overall trend: performance decreased proportionally to the number of missing cell types (Figure S13, Figure S14). This trend was most consistent for the methods that performed best at baseline, particularly cell2location and MuSiC, and was also evident for CARD, RCTD, and SCDC. By contrast, for the methods that already performed relatively poorly at baseline, in particular Seurat and SPOTlight, the deterioration in performance was less pronounced. As observed on the lymph node data, this reflects the fact that baseline performance was already low, leaving less room for further decrease in performance. Note that CARD failed to execute when only two cell types were present in the reference data, and Seurat also failed in this setting. Overall, the results on the hypothalamus data recapitulated the main finding from the lymph node data, namely that deconvolution performance decreases proportionally to the number of cell types missing from the reference data.

**S6.3 Proportions of missing cell types are assigned to transcriptionally similar cell types**

We next examined to which remaining cell types the proportions of removed cell types were reassigned (Figure S15a). To interpret these patterns in the context of transcriptional similarity, we also calculated the pairwise Pearson correlation between the mean expression profiles of the 14 hypothalamic cell types using the selected features (Figure S15b).

For the better-performing methods, reassignment was generally directed toward transcriptionally similar hypothalamic cell types. This was particularly clear for cell2location, MuSiC, CARD, and to a slightly lesser extent RCTD. For example, when upper-layer intratelencephalic cells were removed, their proportions were mainly reassigned to deep-layer intratelencephalic cells, which showed a very high correlation with upper-layer intratelencephalic cells (r = 0.93), and to amygdala excitatory cells (r = 0.94 with upper-layer intratelencephalic). Conversely, when deep-layer intratelencephalic cells were removed, reassignment was mainly directed to upper-layer intratelencephalic (r = 0.93) and corticothalamic and 6b cells (r = 0.91). Similarly, removal of medium spiny neuron cells led primarily to reassignment to eccentric medium spiny neuron cells, and vice versa, consistent with their strong correlation (r = 0.92). Within the oligodendrocyte lineage, removal of oligodendrocyte cells resulted predominantly in reassignment to committed oligodendrocyte precursor cells, which showed the strongest pairwise correlation in the dataset (r = 0.98).

In contrast, lower-performing methods showed more diffuse reassignment patterns, with proportions for the missing cell type distributed across multiple cell types and often accompanied by broader shifts in the predicted proportions of unrelated populations. This is particularly evident for Stereoscope, where the removal of a single cell type affects the proportions of nearly all remaining cell types. Note however, that for the hypothalamus data, in general even the better-performing methods tended to show more diffuse reassignment patterns than observed for the lymph node data, although their reassignment behaviour remained more interpretable overall than that of the lower-performing methods.

We extended our analysis to the removal of two (Figure S16) or three (Figure S17) cell types from the reference data. Overall, the patterns were broadly consistent with those observed in the single-cell-type removal analysis. For the better-performing methods, particularly cell2location, MuSiC, CARD, and RCTD, proportions of missing cell types were often reassigned to transcriptionally similar cell types, although the redistribution was generally more diffuse than with the lymph node dataset. For example, when both medium spiny neuron and eccentric medium spiny neuron cells were removed, reassignment was directed primarily to related neuronal cell types such as splatter and amygdala excitatory, whereas removal of oligodendrocytes and committed oligodendrocyte precursors led mainly to reassignment within the oligodendrocyte lineage (Figure S16). Likewise, when upper-layer intratelencephalic, deep-layer intratelencephalic, and amygdala excitatory were removed together, the proportions of the missing cell types were redistributed predominantly across related excitatory neuronal populations (Figure S17).

**Table S1. Number of cells per cell type in the integrated lymph node single-cell reference dataset**

| **Cell type** | **Number of cells** | **Number of cells**  **(downsampled)** |
| --- | --- | --- |
| Adipocytes | 2711 | 300 |
| Astrocytes | 67 | 0 |
| B cells | 23147 | 300 |
| CD4 T cells | 16441 | 300 |
| CD8 T cells | 6941 | 300 |
| Chondrocytes | 1 | 0 |
| Dendritic cells | 53 | 0 |
| Endothelial cells | 1658 | 300 |
| Epithelial cells | 1 | 0 |
| Erythrocytes | 4 | 0 |
| Fibroblasts | 4720 | 300 |
| Hematopoietic stem cells | 1001 | 300 |
| Macrophages | 1071 | 300 |
| Melanocytes | 2 | 0 |
| Mesangial cells | 1 | 0 |
| Monocytes | 734 | 300 |
| Myocytes | 122 | 122 |
| Neurons | 1 | 0 |
| Neutrophils | 97 | 97 |
| NK cells | 1538 | 300 |
| Skeletal muscle | 569 | 300 |
| Smooth muscle | 3 | 0 |

**Table S2. Cell types grouped by similarity in lymph node dataset**

| **Pairs of similar cell types** | **Triplets of similar cell types** | **Quintuplet of similar cell types** |
| --- | --- | --- |
| CD4 T and CD8 T cells | CD4 T, CD8 T and NK cells | B, CD4 T, CD8 T, NK cells and HSC |
| Myocytes and skeletal muscle | Fibroblasts, myocytes and skeletal muscle |  |
| Hematopoietic stem cells and NK cells | B, CD4 T and CD8 T cells |  |
| Macrophages and monocytes | Macrophages, monocytes and NK cells |  |
| Adipocytes and fibroblasts | Adipocytes, endothelial cells, and fibroblasts |  |

**Table S3. Number of cells per cell type in the hypothalamus single-nucleus reference dataset**

| **Cell type** | **Number of cells** | **Number of cells**  **(downsampled)** |
| --- | --- | --- |
| Amygdala excitatory ^§^ | 1755 | 300 |
| Astrocyte | 1428 | 300 |
| CGE interneuron ^§^ | 1423 | 300 |
| Committed oligodendrocyte precursor | 302 | 300 |
| Deep-layer corticothalamic and 6b ^§^ | 268 | 268 |
| Deep-layer intratelencephalic ^§^ | 571 | 300 |
| Deep-layer near-projecting ^§^ | 156 | 0 |
| Eccentric medium spiny neuron ^§^ | 434 | 300 |
| Ependymal | 1 | 0 |
| Fibroblast | 17 | 0 |
| Hippocampal CA1-3 ^§^ | 1 | 0 |
| LAMP5-LHX6 and Chandelier ^§^ | 286 | 286 |
| Medium spiny neuron ^§^ | 435 | 300 |
| MGE interneuron ^§^ | 1291 | 300 |
| Microglia | 240 | 0 |
| Miscellaneous | 206 | 0 |
| Oligodendrocyte | 307 | 300 |
| Oligodendrocyte precursor | 671 | 300 |
| Splatter ^§^ | 397 | 300 |
| Upper-layer intratelencephalic ^§^ | 2321 | 300 |
| Vascular | 47 | 0 |

^§^ Neuronal cell types

**Table S4. Cell types grouped by similarity in hypothalamus dataset**

| **Pairs of similar cell types** | **Triplets of similar cell types** |
| --- | --- |
| Oligodendrocyte and committed oligodendrocyte precursor | Upper-layer intratelencephalic, deep-layer intratelencephalic, and amygdala excitatory |
| Medium spiny neuron and eccentric medium spiny neuron | Eccentric medium spiny neuron, medium spiny neuron, and splatter |
| Upper-layer intratelencephalic and deep-layer intratelencephalic |  |

**Table S5. Run time per deconvolution method (lymph node data)**

| **Method** | **Run time (mean ± standard deviation, seconds)** |
| --- | --- |
| CARD | 61 ± 0 |
| cell2Location ^§^ | 1023 ± 7 |
| MuSiC | 2303 ± 14 |
| RCTD | 204 ± 3 |
| SCDC | 1333 ± 28 |
| Seurat | 54 ± 5 |
| SPOTlight | 5534 ± 2 |
| Stereoscope ^§^ | 362 ± 2 |

**Table S5.** For each deconvolution method we performed 5 runs with the lymph node reference data and ST1 in the baseline scenario. All methods were run serially to avoid CPU/GPU sharing. Performance was measured on an Ubuntu 22.04.5 machine with 22 CPU cores, 176 GB RAM, dual A10 GPU. Average run time and standard deviation are shown. ^§^ method uses GPU.

**Figure S1. Human lymph node scRNA-seq reference data**

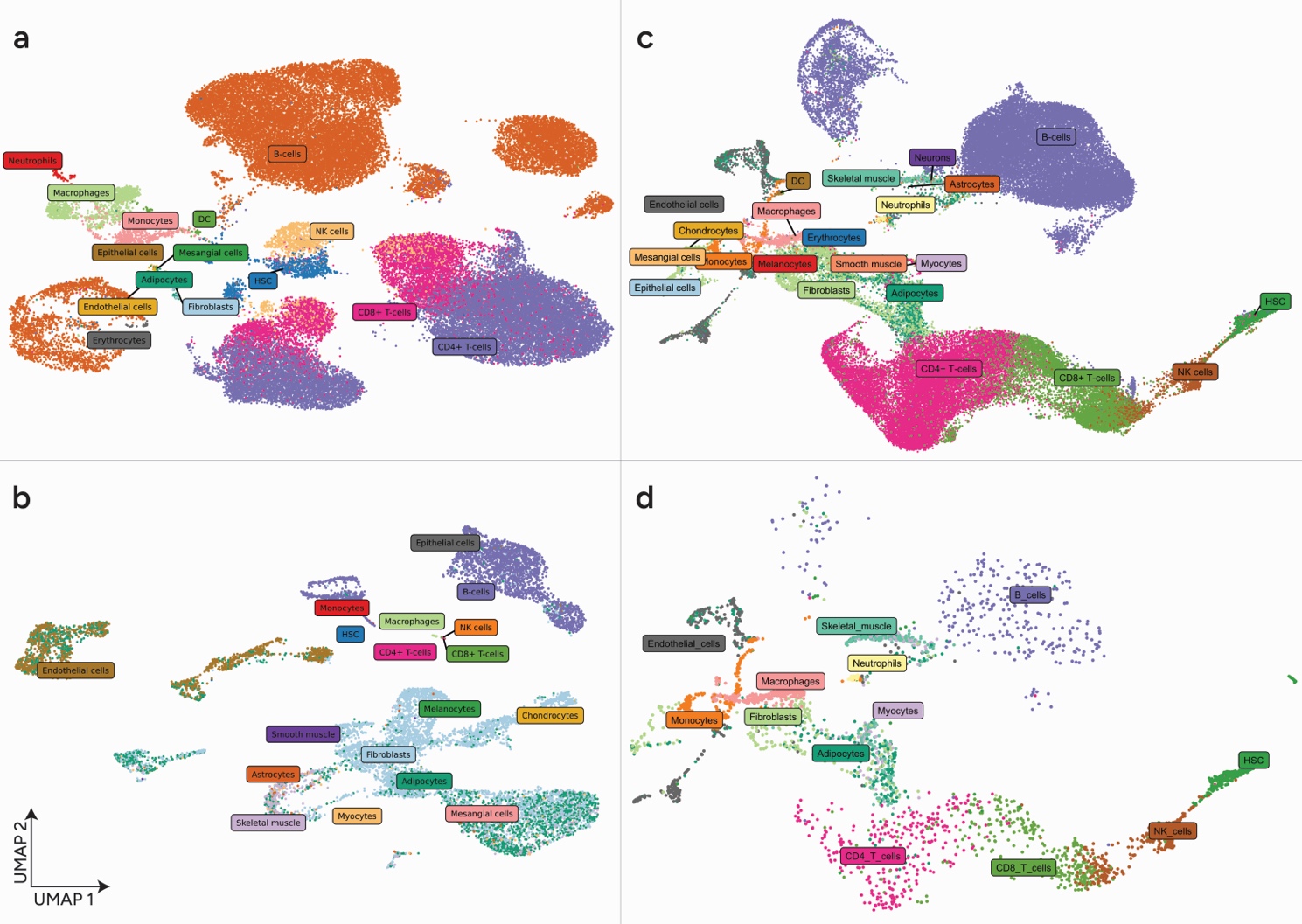

**Figure S1**. **The integrated lymph node single-cell reference dataset captures the major cell populations used for downstream benchmarking.** UMAP representation of human lymph node scRNA-seq datasets used in this study. (a) Tabula Sapiens lymph node data; (b) Lymph node stromal cell data including spiked in B cells; (c) Integrated single-cell data after for the data from (a) and (b) combined and after gene selection; (d) Downsampled integrated dataset. The (counts of the) downsampled data have been used as single-cell reference dataset in downstream analysis.

**Figure S2. Workflow for simulation of spatial transcriptomics data**

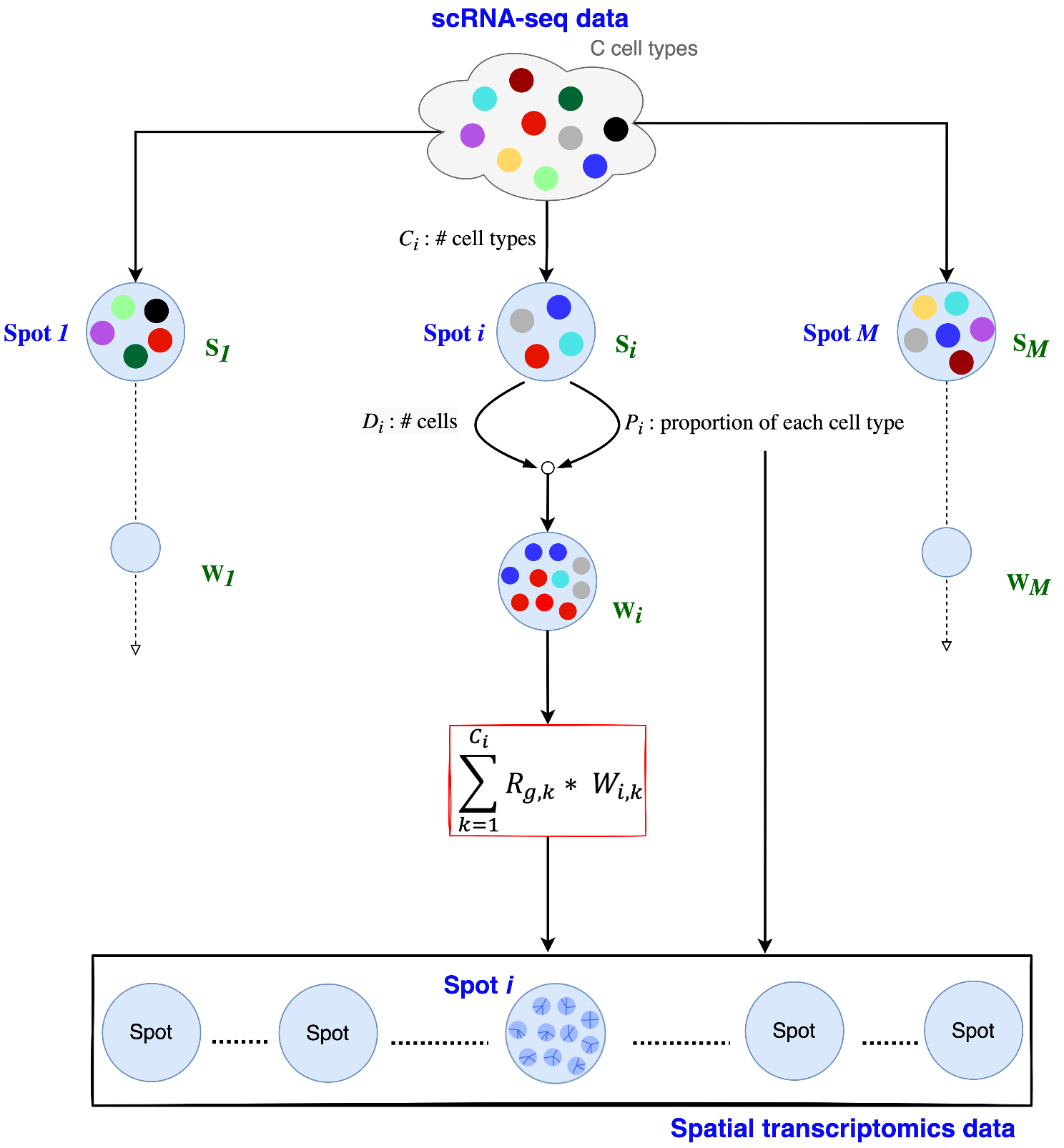

**Figure S2**. **Simulation workflow generates spatial transcriptomics data with known spot-level cell type proportions from a single-cell reference dataset.** Spatial transcriptomics data is simulated from a scRNA-seq reference dataset while varying both the number of cell types C_i_ and the number of cells D_i_ for each spot i. See Section S3 for a detailed description of the different steps of the simulation workflow. In short the notation is as follows, S_i_ denotes the set of cell types present in a spot; P_i,k_ denotes the proportions of cell type k in a spot; W_i,k_ represents the number of cells of cell type k in a spot; R_g,k_ is the mRNA count for gene g in cell type k (in spot i). R_g,k_ is calculated per spot from the counts for gene g of five randomly selected cells of cell type k. Note that since the selected deconvolution methods, with the exception of CARD, do not take spatial correlation into account, spots are treated as independent in our simulation framework.

**Figure S3. Performance at baseline and in case of cell type mismatch for simulated datasets ST2 and ST3 (lymph node data)**

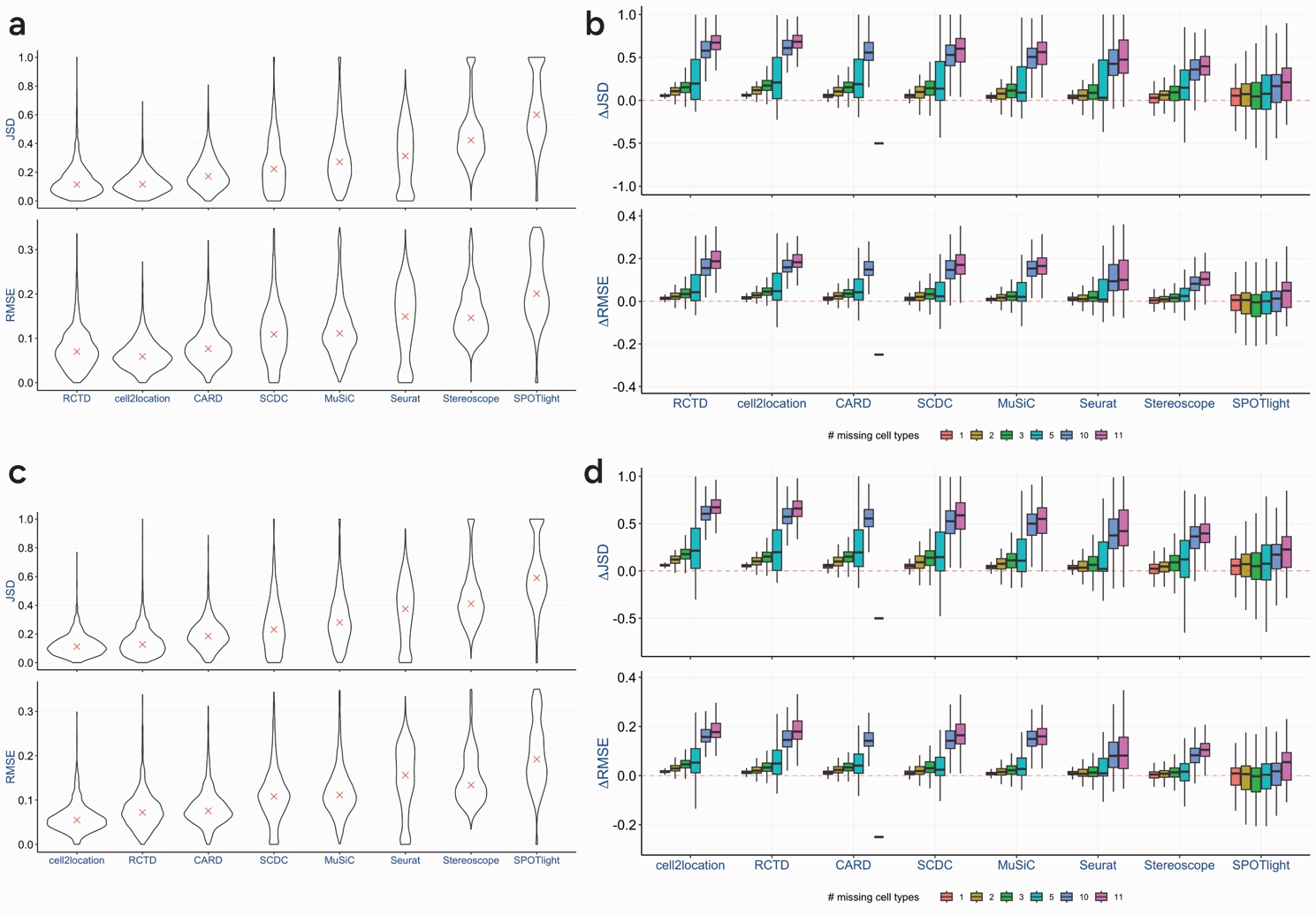

**Figure S3**. **Overall performance trends observed for ST1 are also observed in simulated lymph node datasets ST2 and ST3.** Performance of deconvolution methods for simulated datasets ST2 (a, b) and ST3 (c, d). (a, c) Baseline performance of cell type deconvolution methods. Comparison of the ground truth and predicted proportions using the Jensen-Shannon divergence (JSD) and the root mean square error (RMSE) as performance metrics for simulated dataset ST2 (a) and ST3 (c) for eight cell type deconvolution methods. Violin plots correspond to the distribution of the values of the indicated performance metric across 1,600 spots. Median values are indicated with a red cross. Deconvolution methods in both panels are ordered by increasing median JSD value. (b, d) Performance of cell type deconvolution methods in case of cell type mismatch. Boxplots of the spot-wise difference in performance between the scenario where cell types are missing from the reference data and the baseline scenario (ΔJSD, ΔRMSE) for ST2 (b) and ST3 (d). A value of ΔJSD/ΔRMSE (y-axis) above zero corresponds to a decrease in performance compared to the baseline scenario. The colour key indicates the number of missing cell types (see Section S4, Table S2). ΔJSD/ΔRMSE were calculated as the spot-wise mean across multiple instances of a particular scenario. A dash (-) indicates missing results for that particular scenario. Deconvolution methods in panel (b) and (d) are in the same order as in panel (a) and (c), respectively.

**Figure S4. Per-cell-type RMSE in the baseline scenario for ST1, ST2, and ST3 (lymph node data)**

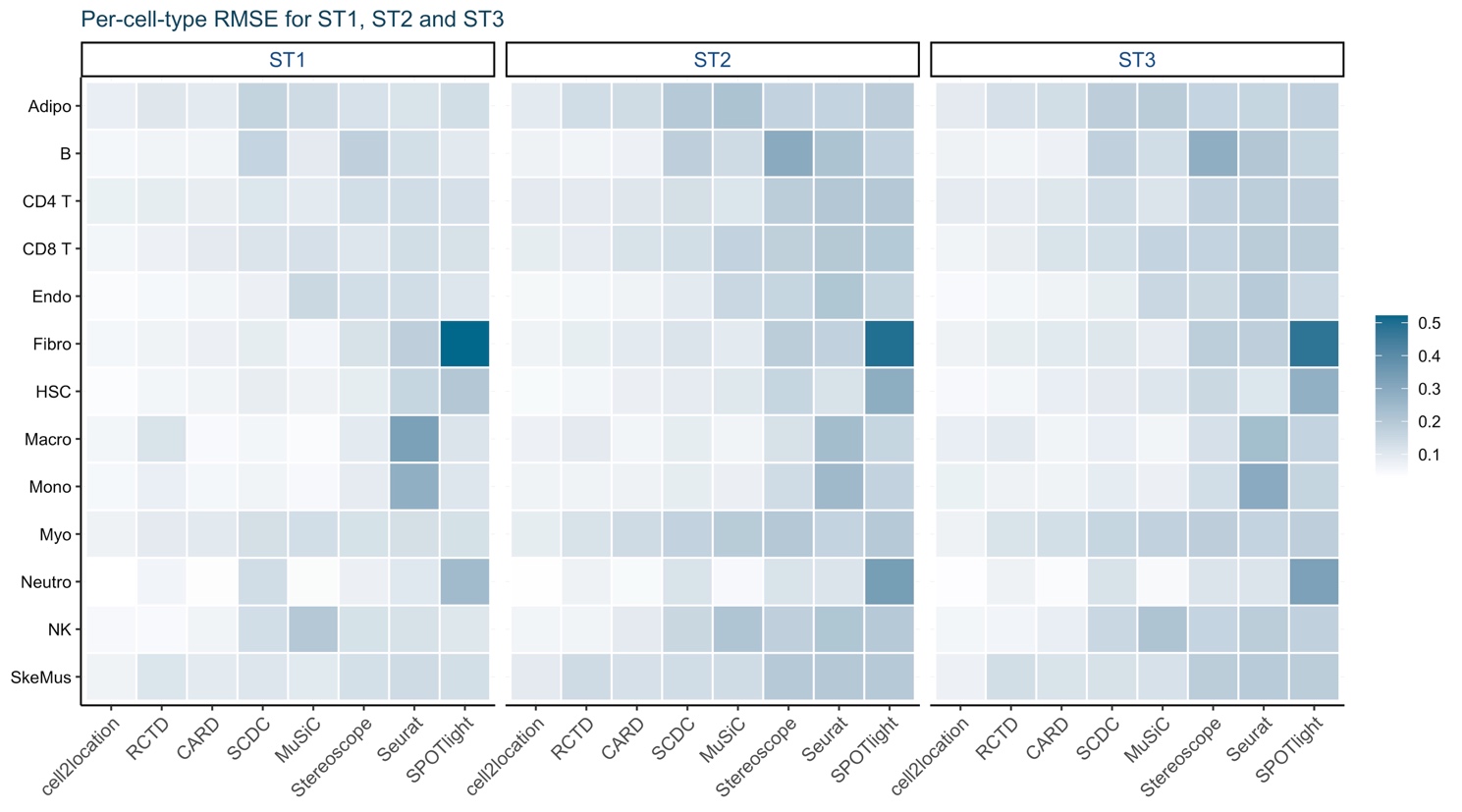

**Figure S4. Baseline deconvolution accuracy varies not only between methods but also between cell types across all three simulated lymph node datasets.** Per-cell-type root mean square error (RMSE) in the baseline scenario for simulated datasets ST1, ST2, and ST3 with lymph node data.

**Figure S5. Performance in case of cell type mismatch (lymph node data)**

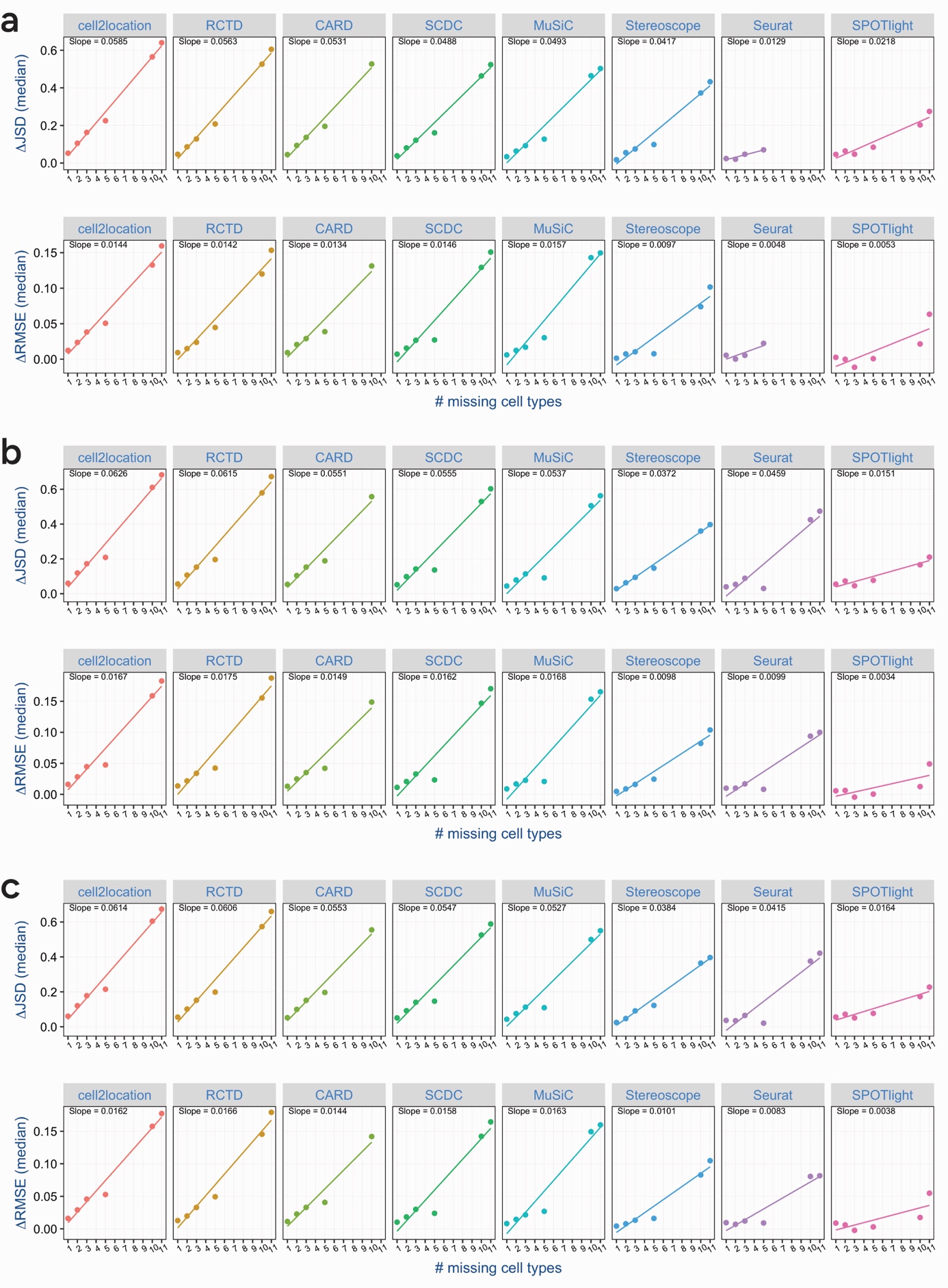

**Figure S5**. **Deconvolution performance decreases proportionally to the number of cell types missing from the lymph node reference data.** Performance in case of cell type mismatch for simulated datasets ST1 (a), ST2 (b), and ST3 (c). (a, b, c) Median values of ΔJSD/ΔRMSE for the different mismatch scenarios per deconvolution method. Linear regression was applied, with fitted lines and slopes displayed. Deconvolution methods are in the same order as in Figure 2.

**Figure S6. Cell type reassignment for removal of one cell type for simulated datasets ST2 and ST3 (lymph node data)**

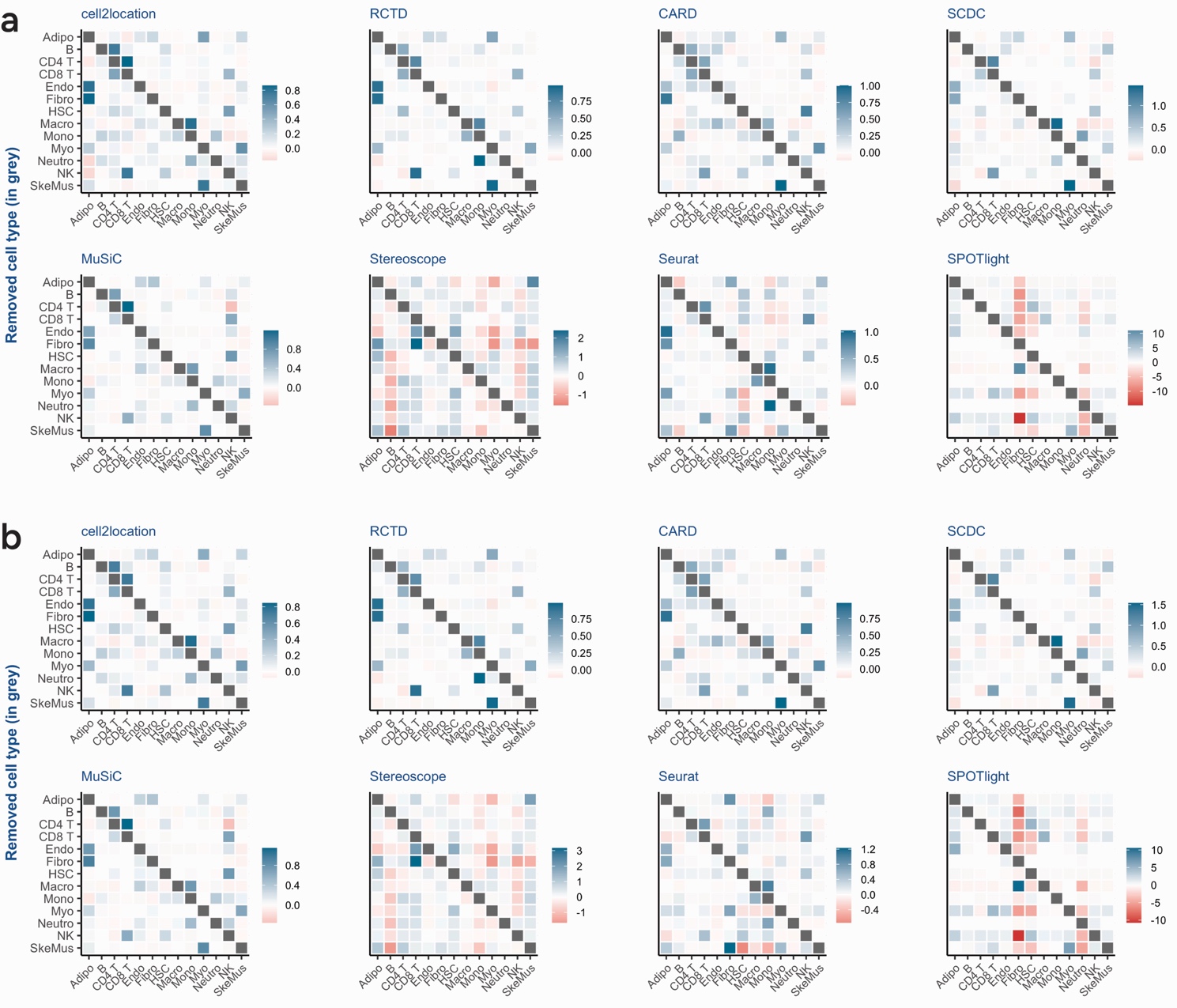

**Figure S6**. **Proportions of cell types missing from the lymph node reference are reassigned to transcriptionally similar cell types.** Cell-type-specific effects for the mismatch scenario with one cell type missing from the lymph node reference data for simulated datasets ST2 (a) and ST3 (b). (a, b) Heatmaps of the reassignment values for each deconvolution method. When a cell type is excluded from the reference data, the reassignment value denotes the normalized change in predicted proportions, relative to the baseline predictions, for each of the remaining cell types (see Section S4). Each row of the heatmap displays the reassignment values for the cell types indicated on the x-axis, following the removal of the cell type indicated on the y-axis (highlighted by the grey cell). Positive values represent an increase in proportion, negative values represent a decrease in proportion (as indicated by the colour key). Deconvolution methods are in the same order as in Figure 2. Adipo: adipocytes; Endo: endothelial cells; Fibro: fibroblasts; HSC: haematopoietic stem cells; Macro: macrophages; Mono: monocytes; Myo: myocytes; Neutro: neutrophils; SkeMus: skeletal muscle cells.

**Figure S7. Cell type reassignment for removal of two cell types (lymph node data)**

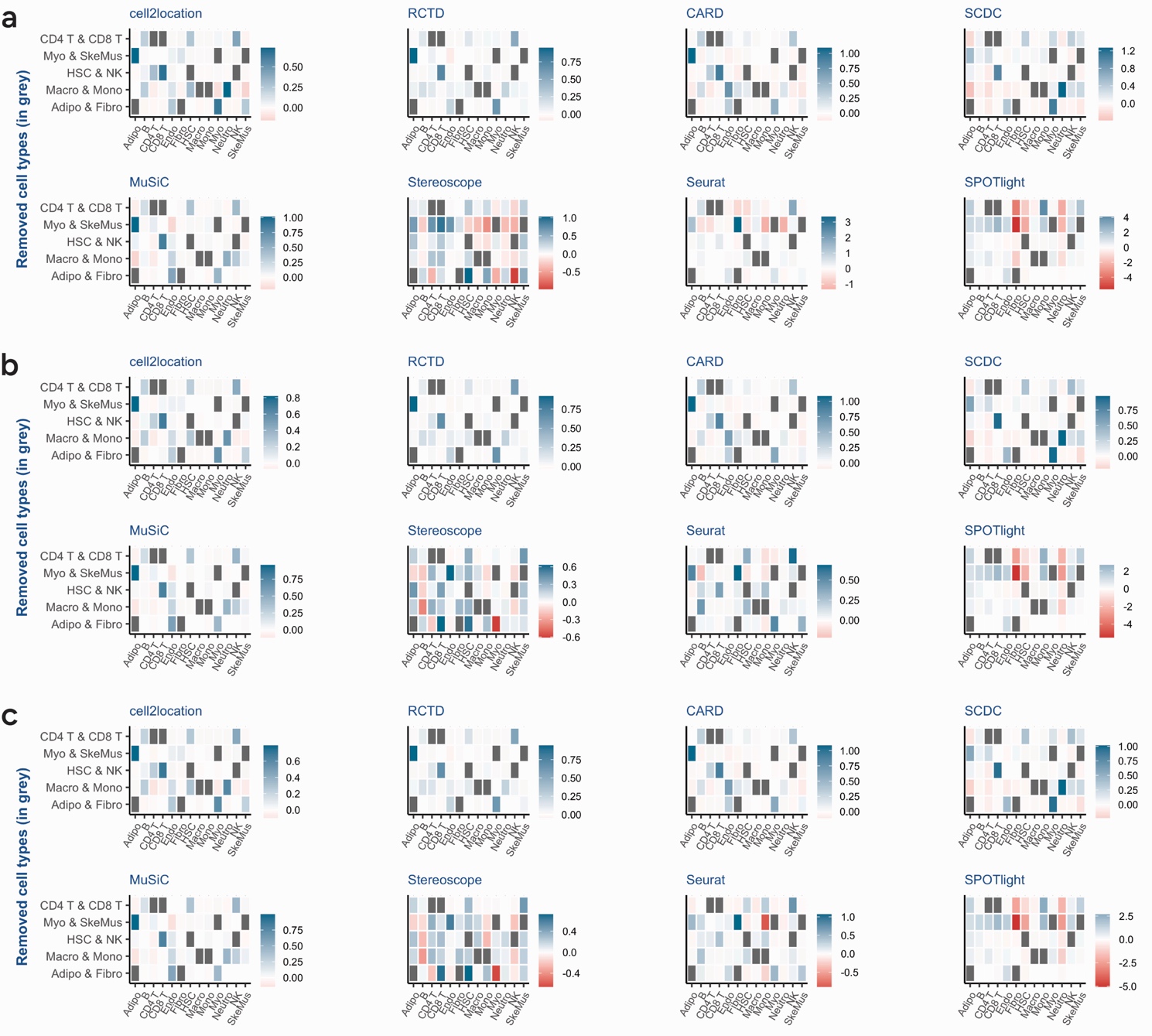

**Figure S7**. **Proportions of cell types missing from the lymph node reference are reassigned to transcriptionally similar cell types.** Cell-type-specific effects for the mismatch scenario with two cell types missing from the reference data for simulated datasets ST1 (a), ST2 (b), and ST3 (c). (a, b, c) Heatmaps of the reassignment values for each deconvolution method. When a pair of cell types is excluded from the reference data, the reassignment value denotes the normalized change in predicted proportions, relative to the baseline predictions, for each of the remaining cell types (see Section S4). Each row of the heatmap displays the reassignment values for the cell types indicated on the x-axis, following the removal of the pair of cell types indicated on the y-axis (highlighted by the grey cells). Positive values represent an increase in proportion, negative values represent a decrease in proportion (as indicated by the colour key). Deconvolution methods are in the same order as in Figure 2. Adipo: adipocytes; Endo: endothelial cells; Fibro: fibroblasts; HSC: haematopoietic stem cells; Macro: macrophages; Mono: monocytes; Myo: myocytes; Neutro: neutrophils; SkeMus: skeletal muscle cells.

**Figure S8. Cell type reassignment for removal of three cell types (lymph node data)**

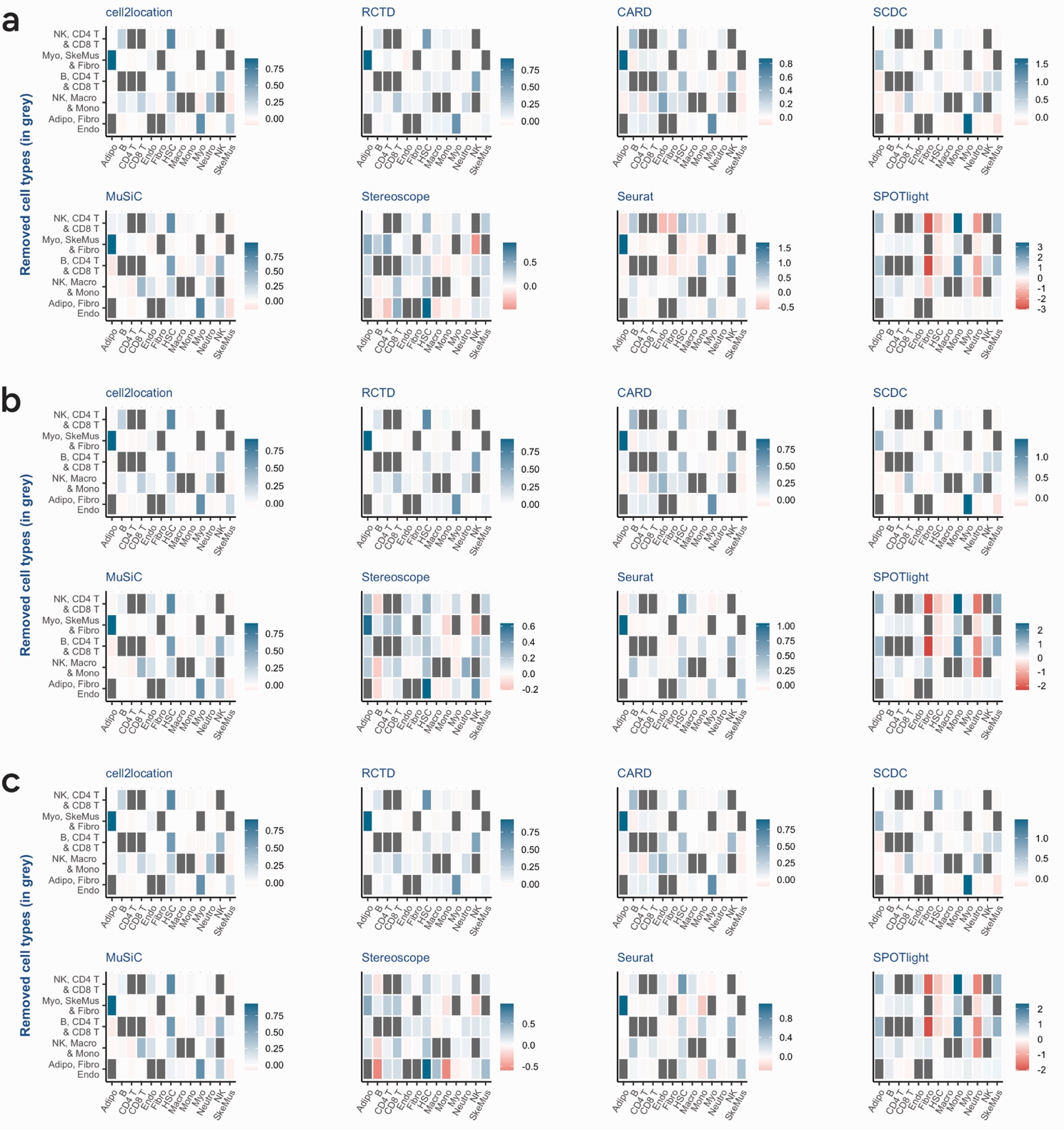

**Figure S8**. **Proportions of cell types missing from the lymph node reference are reassigned to transcriptionally similar cell types.** Cell-type-specific effects for the mismatch scenario with three cell types missing from the reference data for simulated datasets ST1 (a), ST2 (b), and ST3 (c). (a, b, c) Heatmaps of the reassignment values for each deconvolution method. When a triplet of cell types is excluded from the reference data, the reassignment value denotes the normalized change in predicted proportions, relative to the baseline predictions, for each of the remaining cell types (see Section S4). Each row of the heatmap displays the reassignment values for the cell types indicated on the x-axis, following the removal of the triplet of cell types indicated on the y-axis (highlighted by the grey cells). Positive values represent an increase in proportion, negative values represent a decrease in proportion (as indicated by the colour key). Deconvolution methods are in the same order as in Figure 2. Adipo: adipocytes; Endo: endothelial cells; Fibro: fibroblasts; HSC: haematopoietic stem cells; Macro: macrophages; Mono: monocytes; Myo: myocytes; Neutro: neutrophils; SkeMus: skeletal muscle cells.

**Figure S9. Summary of deconvolution results for simulated datasets ST1-3 (lymph node data)**

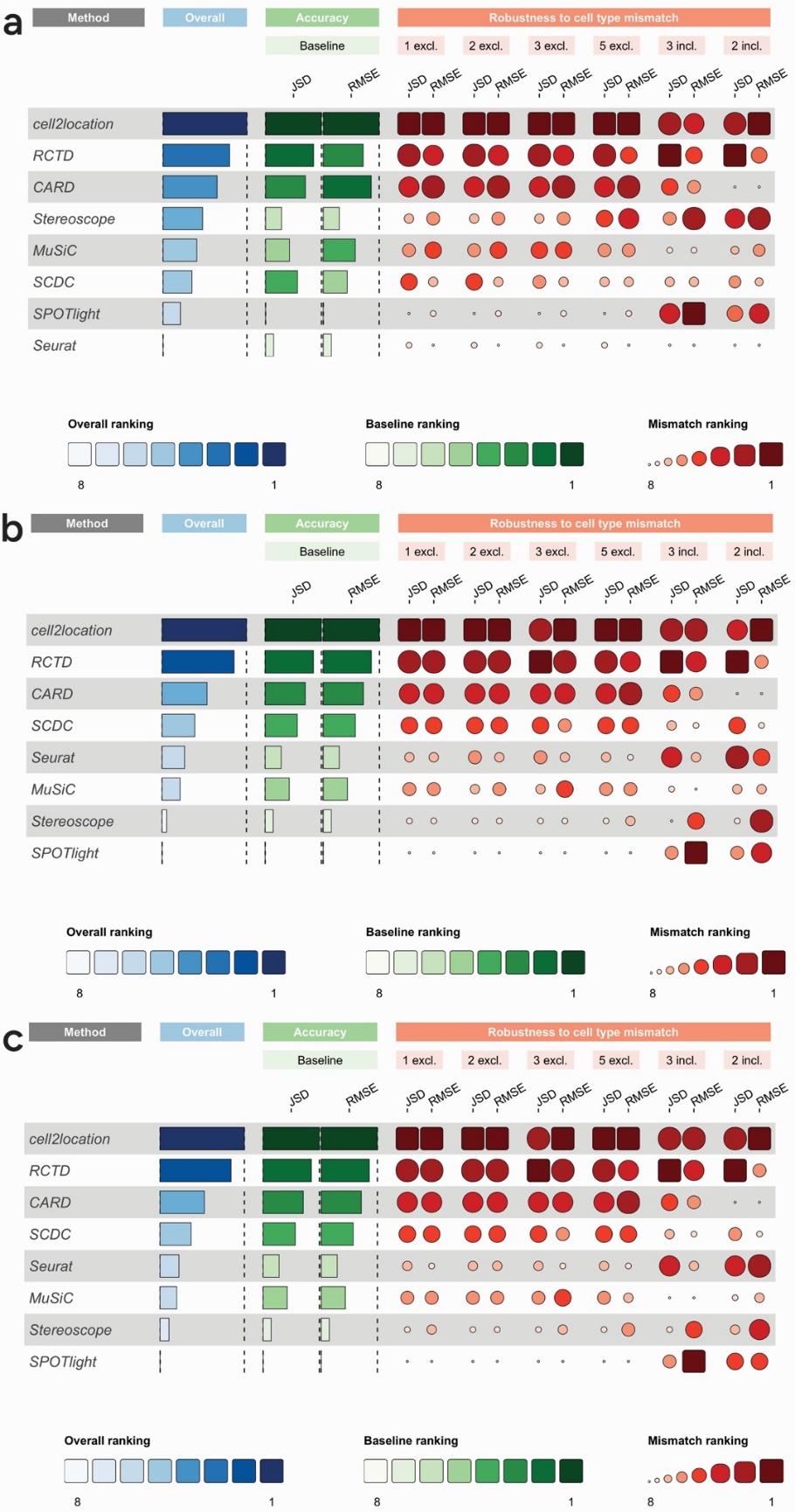

**Figure S9.** **Overall method ranking reflects both baseline accuracy and robustness to cell type mismatch across lymph node simulations.** Summary of deconvolution results for simulated datasets ST1 (a), ST2 (b), and ST3 (c). (a, b, c) Performance is shown at baseline (accuracy; green) and in case of cell type mismatch (robustness to cell type mismatch; red), and summarized as overall performance (blue). Deconvolution methods are ranked based on the mean value of the indicated performance metrics (JSD, RMSE). Darker shades indicate better performance. The overall ranking was computed using the mean of all baseline and mismatch rankings (for both JSD and RMSE)**.** ‘excl.’ indicates the number of cell types missing from the reference dataset, ‘incl.’ indicates the number of cell types remaining in the reference dataset.

**Figure S10. Hypothalamus snRNA-seq reference data**

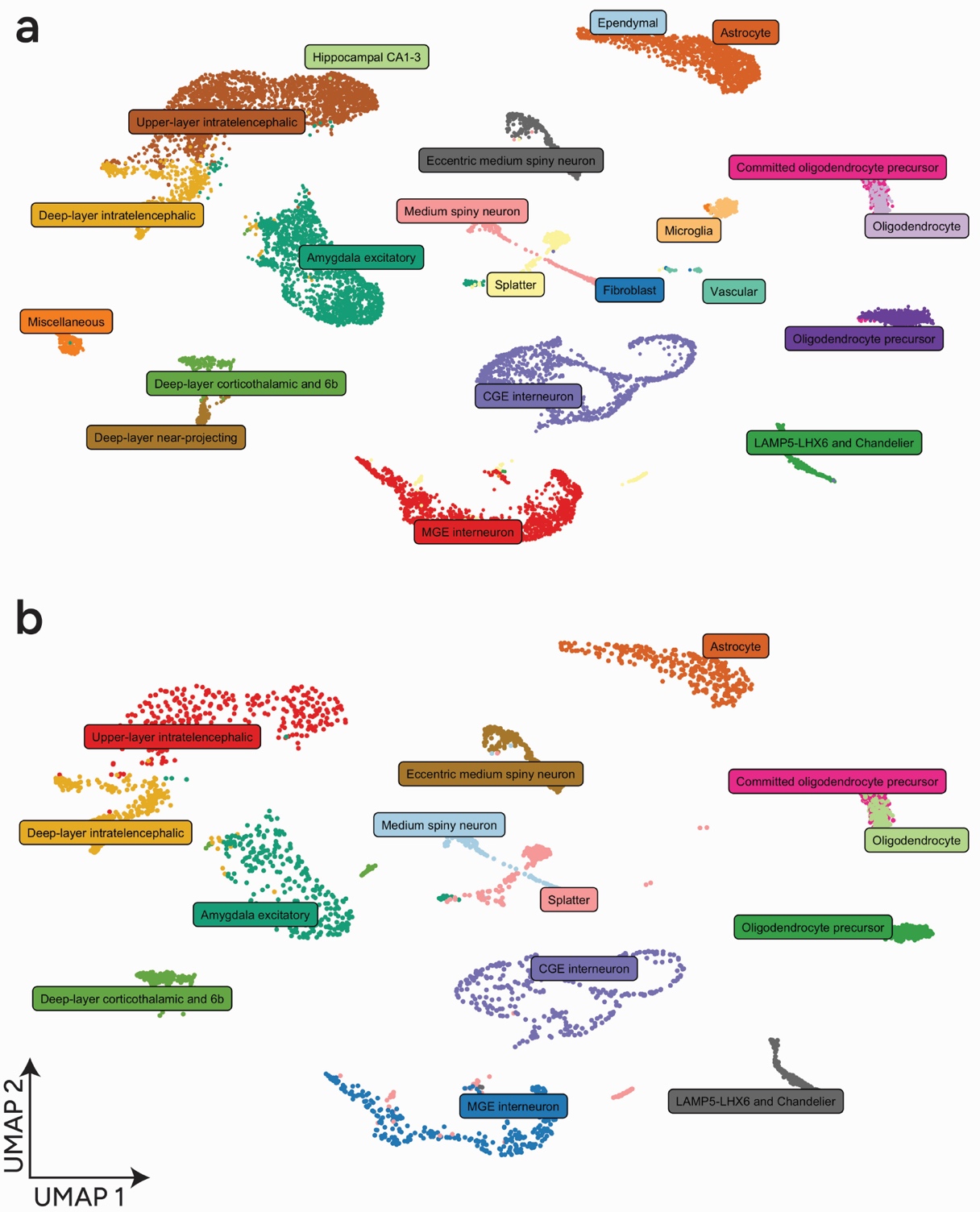

**Figure S10. The processed hypothalamus single-nucleus reference dataset retains the major sufficiently represented cell populations used for benchmarking.** UMAP representation of the human hypothalamus supraoptic region snRNA-seq dataset used in this study. (a) Original dataset containing all cells. (b) Processed and downsampled dataset used as the single-cell reference dataset in downstream analysis after retaining sufficiently represented cell types and downsampling overrepresented populations.

**Figure S11. Performance at baseline for simulated dataset ST1 (hypothalamus data)**

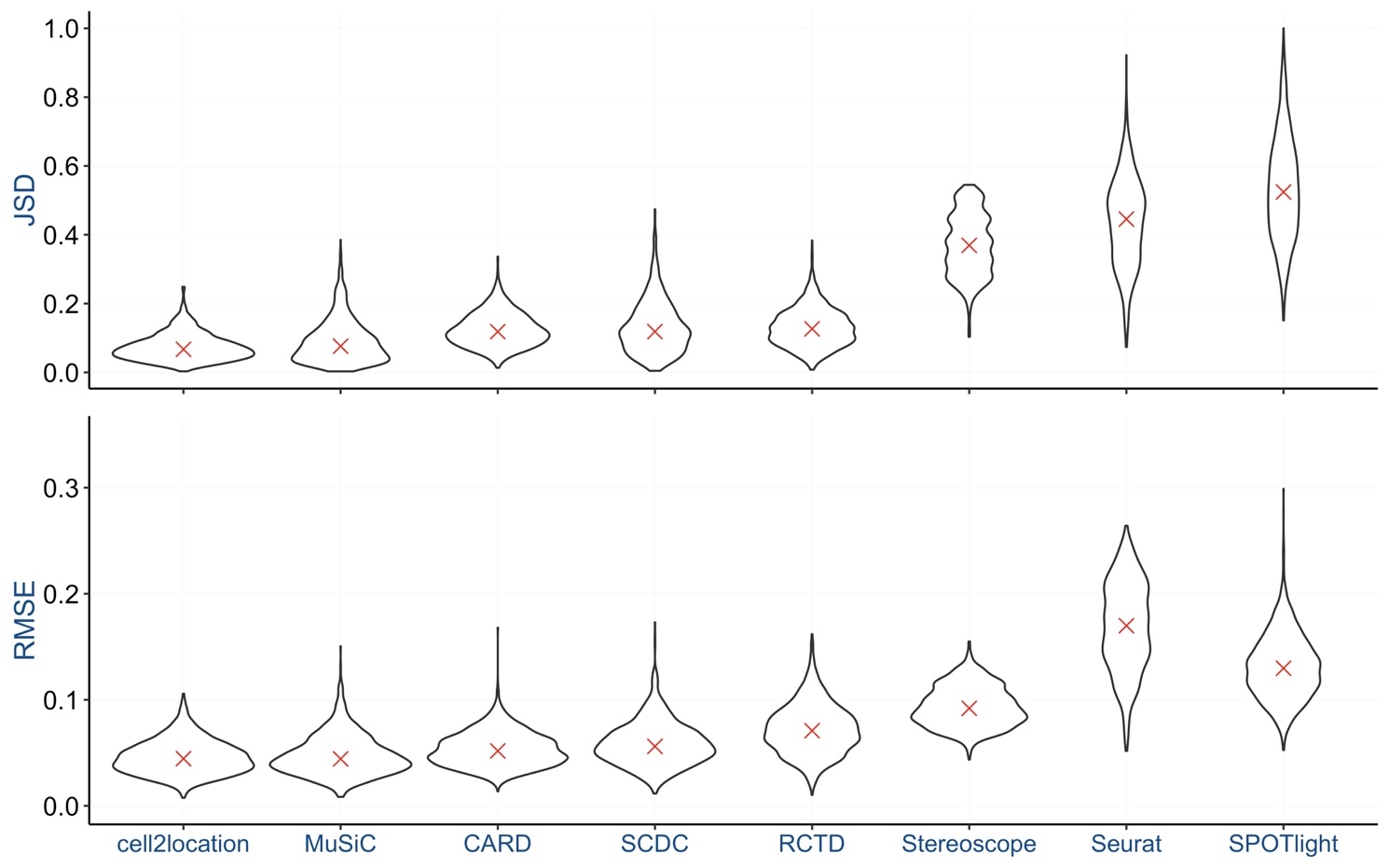

**Figure S11. Clear performance differences between deconvolution methods are observed at baseline for the hypothalamus data.** Comparison of the ground truth and predicted proportions using the Jensen-Shannon divergence (JSD) and the root mean square error (RMSE) as performance metrics for eight cell type deconvolution methods. Violin plots correspond to the distribution of the values of the indicated performance metric across 1,600 spots. Median values are indicated with a red cross. Deconvolution methods are ordered by increasing median JSD value.

**Figure S12. Per-cell-type RMSE in the baseline scenario for ST1 (hypothalamus data)**

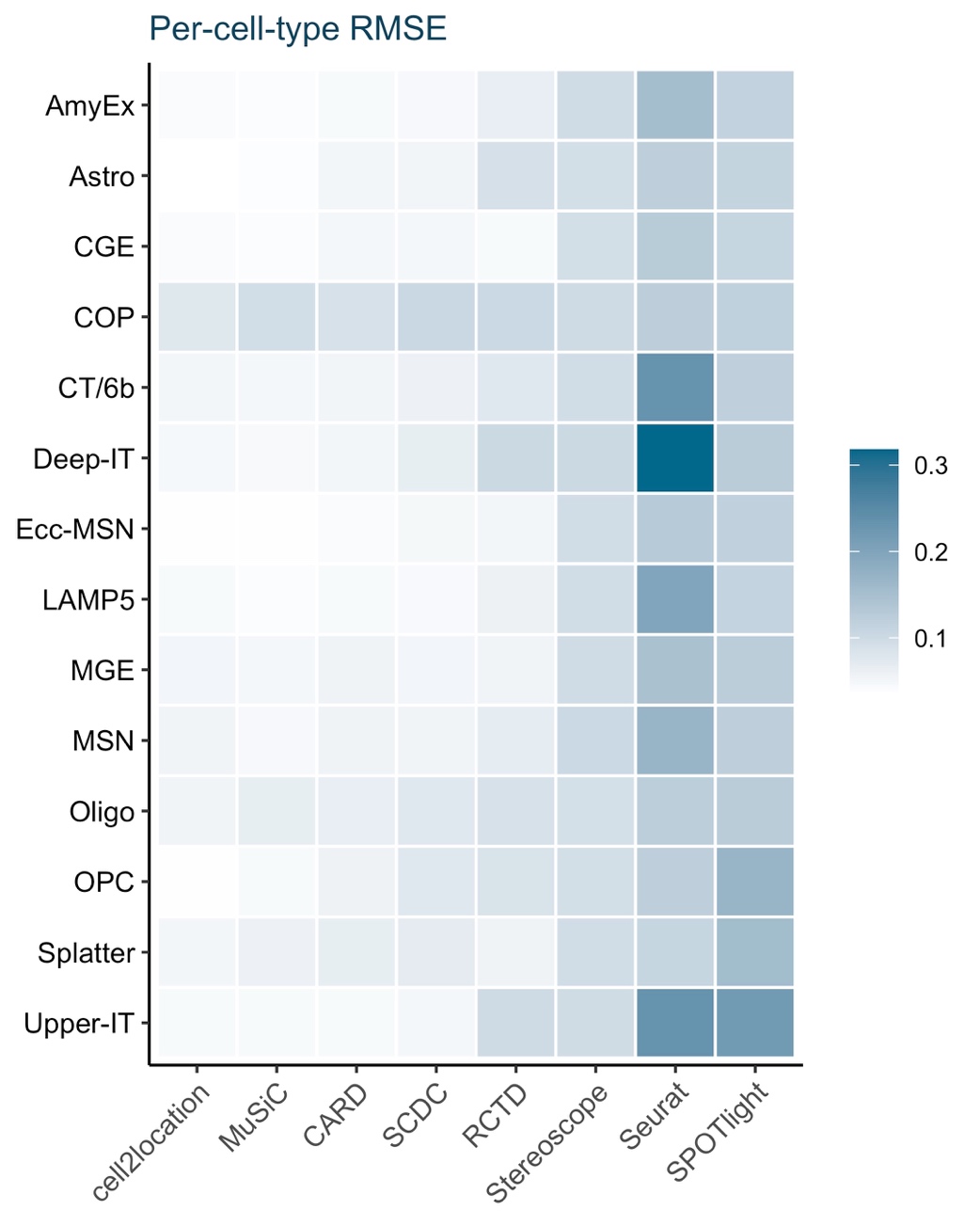

**Figure S12. Baseline deconvolution accuracy varies not only between methods but also between cell types with hypothalamus data.** Per-cell-type root mean square error (RMSE) in the baseline scenario for simulated dataset ST1 with hypothalamus data. Deconvolution methods are in the same order as in Figure S11. AmyEx: Amygdala excitatory; Astro: Astrocyte; CGE: CGE interneuron; COP: Committed oligodendrocyte precursor; CT/6b: Deep-layer corticothalamic and 6b; Deep-IT: Deep-layer intratelencephalic; Ecc-MSN: Eccentric medium spiny neuron; LAMP5: LAMP5-LHX6 and Chandelier; MGE: MGE interneuron; MSN: Medium spiny neuron; Oligo: Oligodendrocyte; OPC: Oligodendrocyte precursor; Upper-IT: Upper-layer intratelencephalic.

**Figure S13. Performance in case of cell type mismatch (hypothalamus data)**

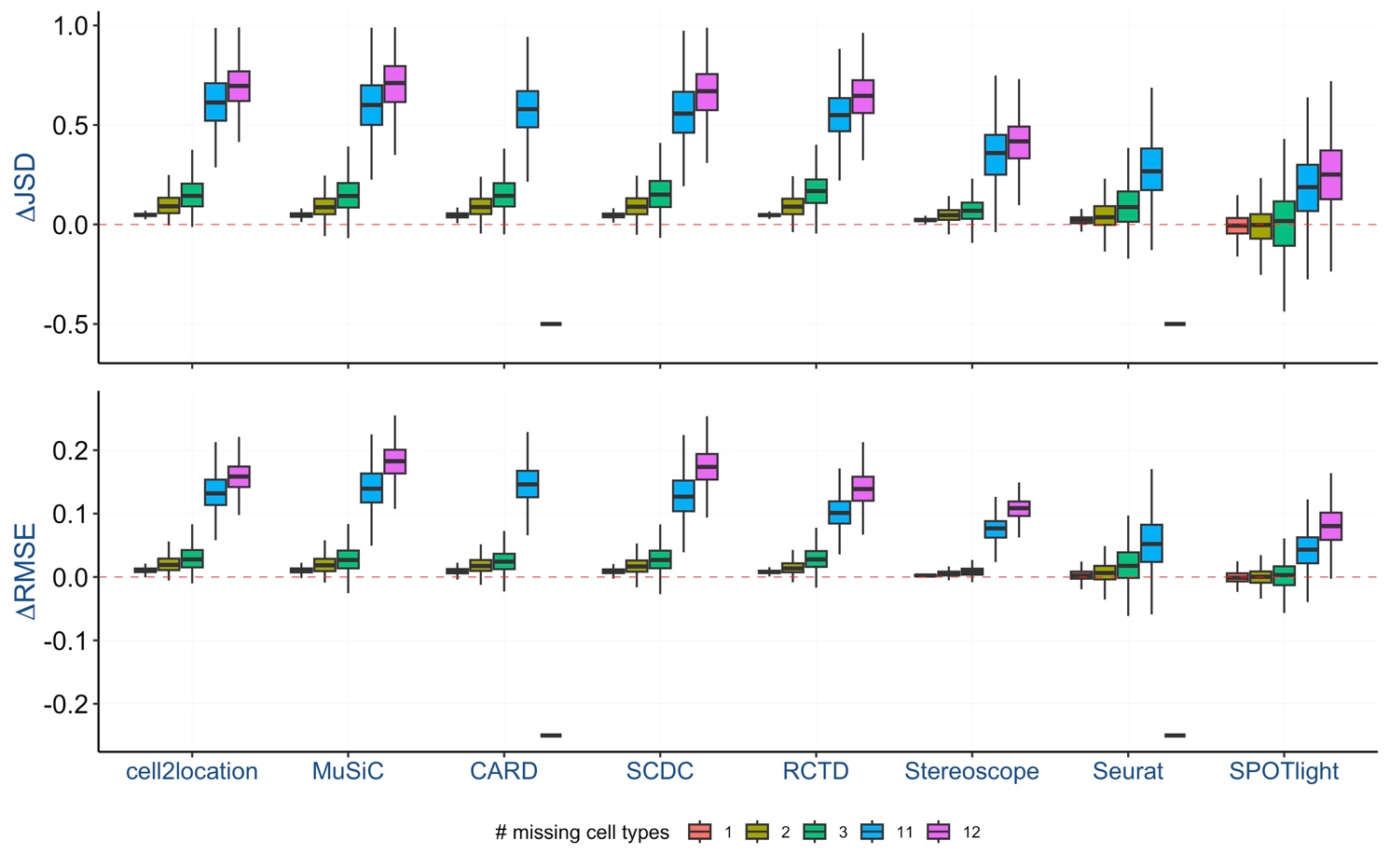

**Figure S13. Performance worsens when cell types are missing from the hypothalamus reference data.** Performance of cell type deconvolution methods in case of cell type mismatch for simulated dataset ST1 with hypothalamus data. Boxplots show the spot-wise difference in performance between the scenario where cell types are missing from the reference data and the baseline scenario (ΔJSD, ΔRMSE). A value of ΔJSD/ΔRMSE above zero corresponds to a decrease in performance compared to the baseline scenario. The colour key indicates the number of missing cell types (see Section S5, Table S4). ΔJSD/ΔRMSE were calculated as the spot-wise mean across multiple instances of a particular scenario. A dash (-) indicates missing results for that particular scenario. Deconvolution methods are in the same order as in Figure S11.

**Figure S14. Performance in case of cell type mismatch (hypothalamus data)**

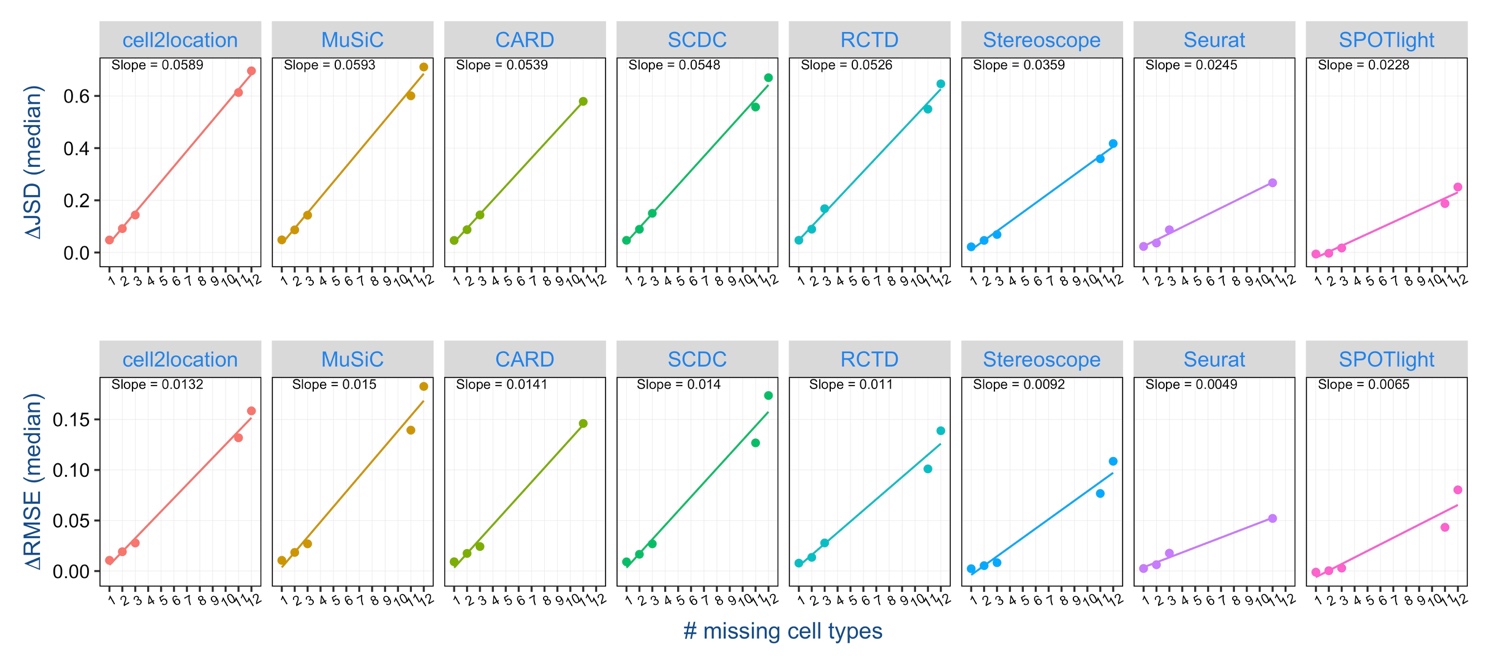

**Figure S14. Deconvolution performance decreases proportionally to number of cell types missing from the hypothalamus reference data.** Median values of ΔJSD and ΔRMSE for the different mismatch scenarios per deconvolution method for simulated dataset ST1 are shown. Linear regression was applied, with fitted lines and slopes displayed. Deconvolution methods are in the same order as in Figure S11.

**Figure S15. Cell type reassignment for removal of one cell type for simulated dataset ST1 (hypothalamus data)**

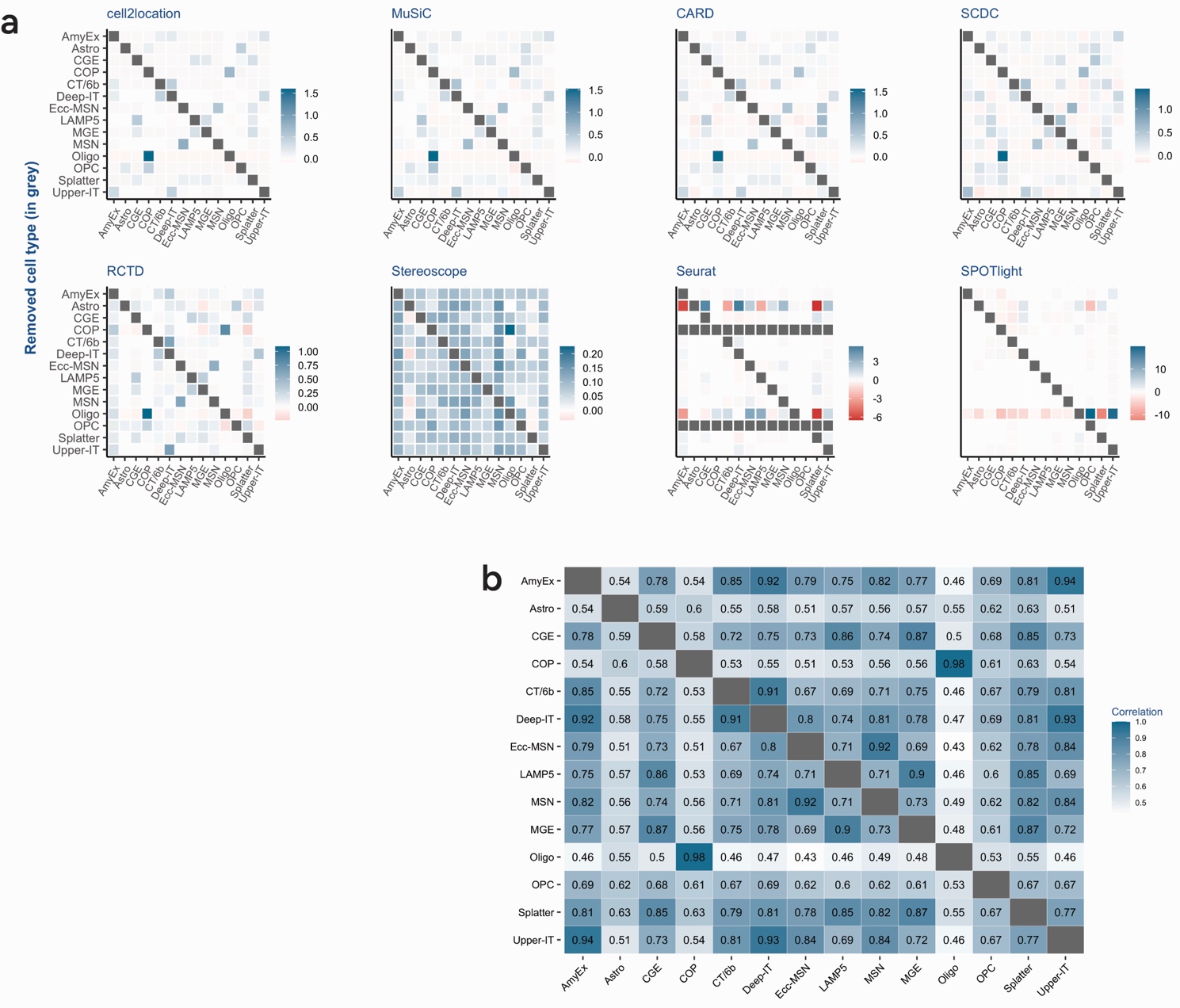

**Figure S15. Proportions of cell types missing from the hypothalamus reference are reassigned to transcriptionally similar cell types.** (a) Heatmaps of the reassignment values for each deconvolution method. When a cell type is excluded from the reference data, the reassignment value denotes the normalized change in predicted proportions, relative to the baseline predictions, for each of the remaining cell types (see Section S5). Each row of the heatmap displays the reassignment values for the cell types indicated on the x-axis, following the removal of the cell type indicated on the y-axis, highlighted by the grey cell. Positive values represent an increase in proportion and negative values represent a decrease in proportion, as indicated by the colour key. For Seurat, all cells are grey for COP and Oligo, since their predicted proportions at baseline are zero for all spots, in which case the reassignment value is undefined. Deconvolution methods are in the same order as in Figure S11. (b) Pairwise Pearson correlation between the mean expression profiles of the 14 hypothalamic cell types using the selected features. AmyEx: Amygdala excitatory; Astro: Astrocyte; CGE: CGE interneuron; COP: Committed oligodendrocyte precursor; CT/6b: Deep-layer corticothalamic and 6b; Deep-IT: Deep-layer intratelencephalic; Ecc-MSN: Eccentric medium spiny neuron; LAMP5: LAMP5-LHX6 and Chandelier; MGE: MGE interneuron; MSN: Medium spiny neuron; Oligo: Oligodendrocyte; OPC: Oligodendrocyte precursor; Upper-IT: Upper-layer intratelencephalic.

**Figure S16. Cell type reassignment for removal of two cell types (hypothalamus data)**

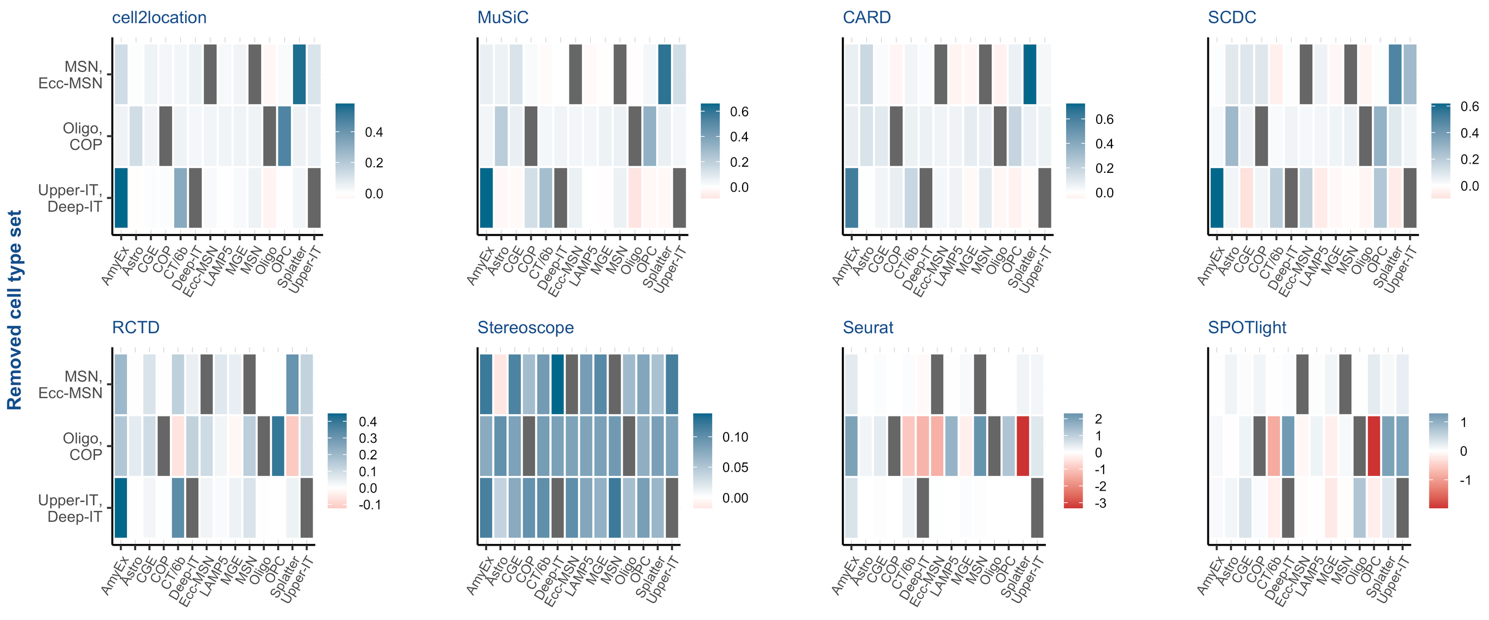

**Figure S16. Proportions of cell types missing from the hypothalamus reference are reassigned to transcriptionally similar cell types.** Heatmaps of the reassignment values for each deconvolution method are shown. When a pair of cell types is excluded from the reference data, the reassignment value denotes the normalized change in predicted proportions, relative to the baseline predictions, for each of the remaining cell types (see Section S5). Each row of the heatmap displays the reassignment values for the cell types indicated on the x-axis, following the removal of the pair of cell types indicated on the y-axis, highlighted by the grey cells. Positive values represent an increase in proportion and negative values represent a decrease in proportion, as indicated by the colour key. Deconvolution methods are in the same order as in Figure S11. AmyEx: Amygdala excitatory; Astro: Astrocyte; CGE: CGE interneuron; COP: Committed oligodendrocyte precursor; CT/6b: Deep-layer corticothalamic and 6b; Deep-IT: Deep-layer intratelencephalic; Ecc-MSN: Eccentric medium spiny neuron; LAMP5: LAMP5-LHX6 and Chandelier; MGE: MGE interneuron; MSN: Medium spiny neuron; Oligo: Oligodendrocyte; OPC: Oligodendrocyte precursor; Upper-IT: Upper-layer intratelencephalic.

**Figure S17. Cell type reassignment for removal of three cell types (hypothalamus data)**

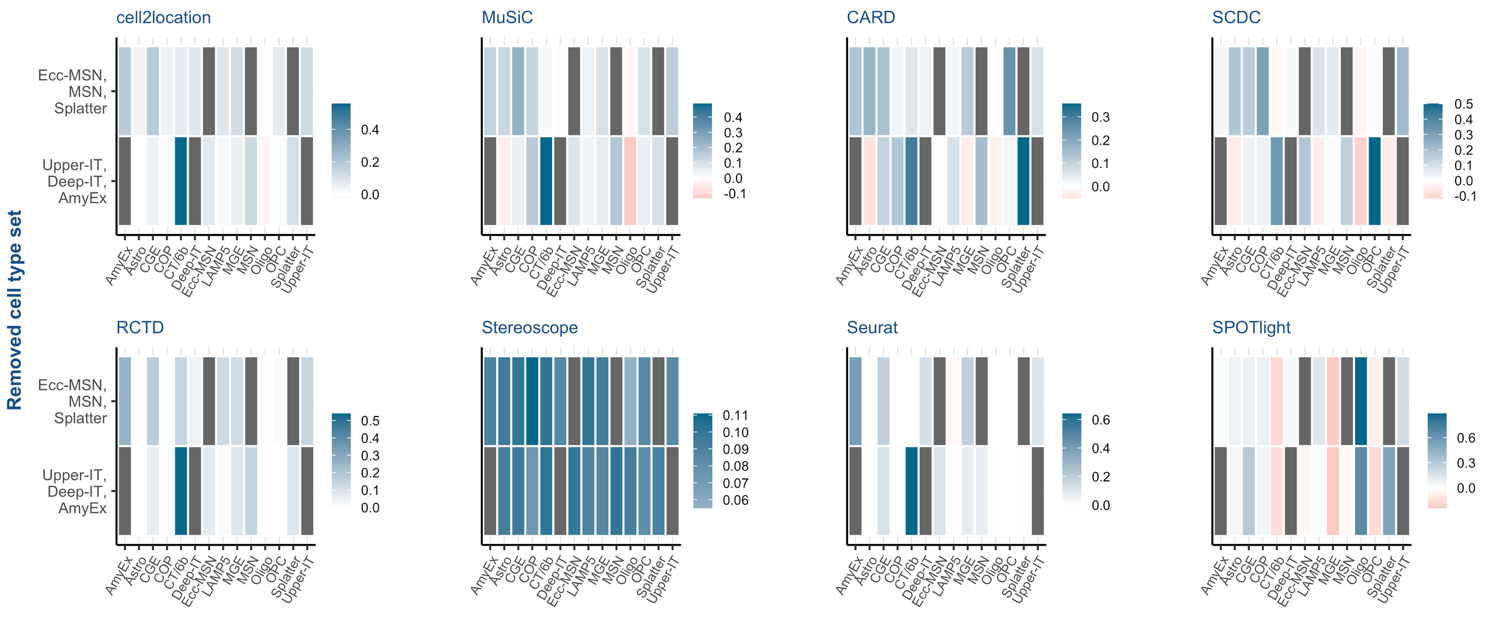

**Figure S17. Proportions of cell types missing from the hypothalamus reference are reassigned to transcriptionally similar cell types.** Heatmaps of the reassignment values for each deconvolution method are shown. When a triplet of cell types is excluded from the reference data, the reassignment value denotes the normalized change in predicted proportions, relative to the baseline predictions, for each of the remaining cell types (see Section S5). Each row of the heatmap displays the reassignment values for the cell types indicated on the x-axis, following the removal of the triplet of cell types indicated on the y-axis, highlighted by the grey cells. Positive values represent an increase in proportion and negative values represent a decrease in proportion, as indicated by the colour key. Deconvolution methods are in the same order as in Figure S11. AmyEx: Amygdala excitatory; Astro: Astrocyte; CGE: CGE interneuron; COP: Committed oligodendrocyte precursor; CT/6b: Deep-layer corticothalamic and 6b; Deep-IT: Deep-layer intratelencephalic; Ecc-MSN: Eccentric medium spiny neuron; LAMP5: LAMP5-LHX6 and Chandelier; MGE: MGE interneuron; MSN: Medium spiny neuron; Oligo: Oligodendrocyte; OPC: Oligodendrocyte precursor; Upper-IT: Upper-layer intratelencephalic.

**Figure S18. Summary of deconvolution results for simulated dataset ST1 (hypothalamus data)**

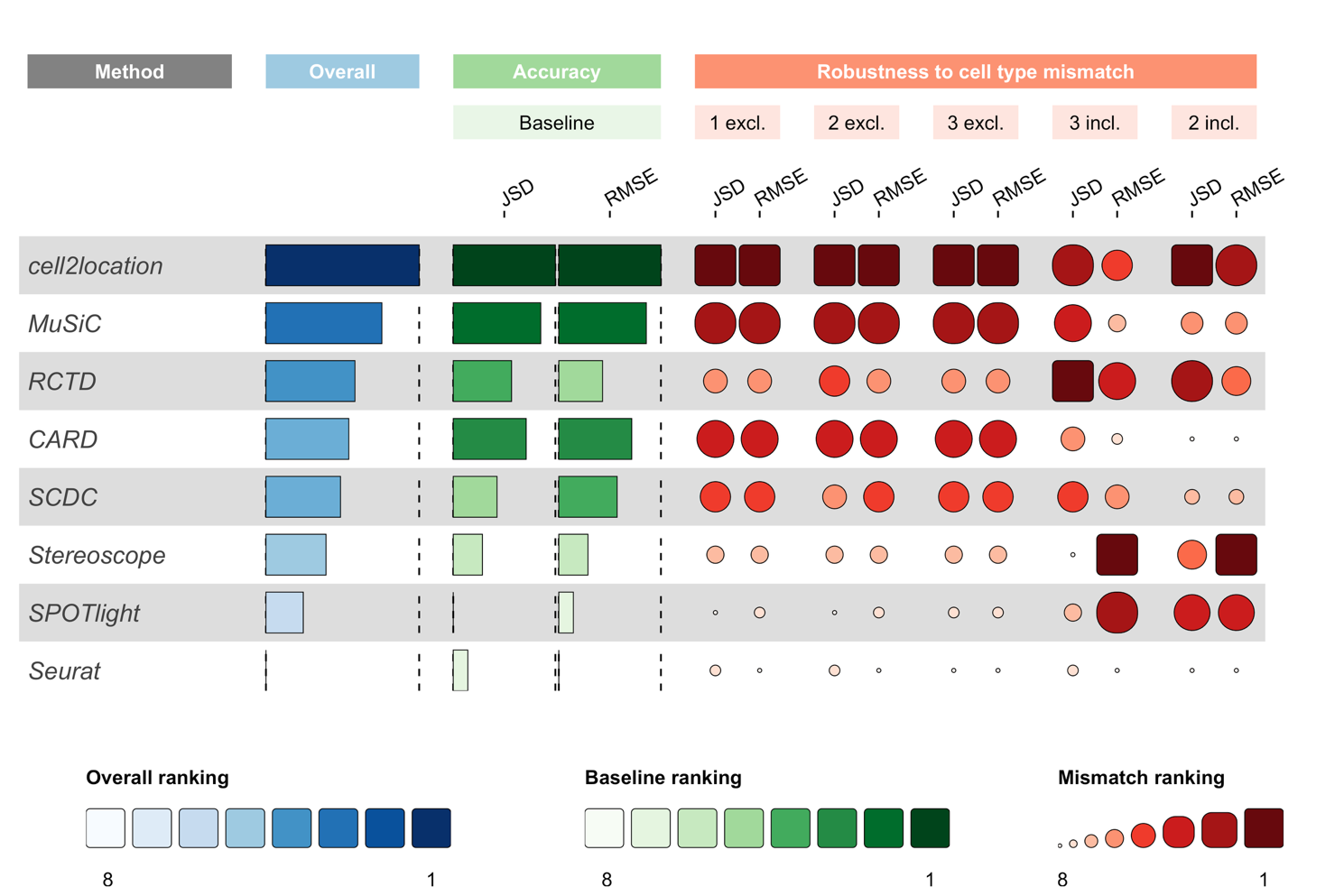

**Figure S18. Overall method ranking reflects both baseline accuracy and robustness to cell type mismatch across hypothalamus simulations.** Summary of deconvolution results for simulated dataset ST1 with hypothalamus data. Performance is shown at baseline (accuracy; green) and in case of cell type mismatch (robustness to cell type mismatch; red), and summarized as overall performance (blue). Deconvolution methods are ranked based on the mean value of the indicated performance metrics (JSD, RMSE). Darker shades indicate better performance. The overall ranking was computed using the mean of all baseline and mismatch rankings for both JSD and RMSE. ‘excl.’ indicates the number of cell types missing from the reference dataset, ‘incl.’ indicates the number of cell types remaining in the reference dataset.
